# Constitutive overexpression of RAM1 increases arbuscule density during arbuscular mycorrhizal symbiosis in Brachypodium distachyon

**DOI:** 10.1101/2020.06.15.146233

**Authors:** Lena M. Müller, Lidia Campos-Soriano, Veronique Levesque-Tremblay, Armando Bravo, Dierdra A. Daniels, Sunita Pathak, Hee-Jin Park, Maria J. Harrison

## Abstract

Arbuscular mycorrhizal (AM) symbiosis is a mutually beneficial association of plants and fungi of the sub-phylum Glomeromycotina. The endosymbiotic AM fungi colonize the inner cortical cells of the roots, where they form branched hyphae called arbuscules that function in nutrient exchange with the plant. To support arbuscule development and subsequently bidirectional nutrient exchange, the root cortical cells undergo substantial transcriptional re-programming. *REDUCED ARBUSCULAR MYCORRHIZA 1 (RAM1)*, studied in several dicot plant species, is a major regulator of this cortical cell transcriptional program. Here, we generated *ram1* mutants and *RAM1* overexpressors in a monocot, *Brachypodium distachyon*. The AM phenotypes of two *ram1* line*s* revealed that *RAM1* is only partly required to enable arbuscule development in *B. distachyon*. Transgenic lines constitutively overexpressing *BdRAM1* showed constitutive expression of AM-inducible genes even in the shoots. Following inoculation with AM fungi, *BdRAM1*-overexpressing roots showed higher arbuscule densities relative to controls, indicating the potential to manipulate the relative proportion of symbiotic interfaces via modulation of *RAM1*. However, the overexpressors also show altered expression of hormone biosynthesis genes and aberrant growth patterns including stunted bushy shoots and poor seed set. While these phenotypes possibly provide additional clues about *BdRAM1*’s scope of influence, they also indicate that directed approaches to increase the density of symbiotic interfaces will require a more focused, potentially cell-type specific manipulation of transcription factor gene expression.

## Introduction

The GRAS (for GA3 INSENSITIVE [GAI], REPRESSOR OF GAI [RGA], and SCARECROW [SCR]) transcription factor *REDUCED ARBUSCULAR MYCORRHIZA (RAM1)* has been characterized in three dicot plant species where it is a major regulator of AM symbiosis. In *Medicago truncatula, Lotus japonicus*, and *Petunia hybrida ram1* mutants, AM fungi display limited arbuscule branching and reduced hyphal colonization of the root, which results in a non-functional symbiosis (Gobbato et al., 2013; Park et al., 2015; Rich et al., 2015; Xue et al., 2015; Pimprikar et al., 2016). *RAM1* expression is induced in colonized cortical cells and is regulated by CYCLOPS, a transcription factor of the common symbiosis signaling pathway (Pimprikar et al., 2016) and also by DELLA proteins (Park et al., 2015; Pimprikar et al., 2016), negative regulators of GA signaling (Daviere and Achard, 2013). RNA sequencing of *ram1* mutants (Luginbuehl et al., 2017) as well as smaller scale gene expression analyses of roots overexpressing *RAM1* (Park et al., 2015; Jiang et al., 2017), indicate that RAM1 either directly or indirectly regulates expression of several symbiosis-associated transcription factors, including the GRAS transcription factor *RAD1*, and three AP2-domain transcription factors of the *WRINKLED5 (WRI5)* family. RAM1 also either directly or indirectly regulates expression of genes involved in the production and transfer of lipids to the fungal symbiont (e.g. *FatM, RAM2, STR*) and the phosphate transporter *PT4* (Gobbato et al., 2012; Park et al., 2015; Pimprikar et al., 2016; Jiang et al., 2017; Luginbuehl et al., 2017). However, so far, only one lipid biosynthesis gene, *RAM2*, has been established as a direct target of RAM1 (Gobbato et al., 2012). Regulation of the other lipid biosynthesis and transport genes likely occurs indirectly through the action of the *WRI5* family genes (Luginbuehl et al., 2017; Jiang et al., 2018).

RAD1, a GRAS transcription factor very closely related to RAM1 (Supplemental Fig. 1) (Park et al., 2015; Xue et al., 2015) is also required for AM symbiosis. In *L. japonicus rad1* mutants, AM fungi display defective arbuscule branching phenotypes reminiscent of those seen in *ram1* (Xue et al., 2015); however, in *M. truncatula rad1*, AM fungi show normal arbuscule branching but reduced colonization levels (Park et al., 2015). In line with this observation, several predicted *RAM1* target genes were induced in colonized *L. japonicus ram1* mutants but induction was completely abolished in *M. truncatula* and *P. hybrida ram1* (Park et al., 2015; Rich et al., 2015; Pimprikar et al., 2016; Luginbuehl et al., 2017). Thus, there are slight differences in regulation of AM symbiosis genes even between relatively closely related plant species (Pimprikar and Gutjahr, 2018).

**Figure 1:**
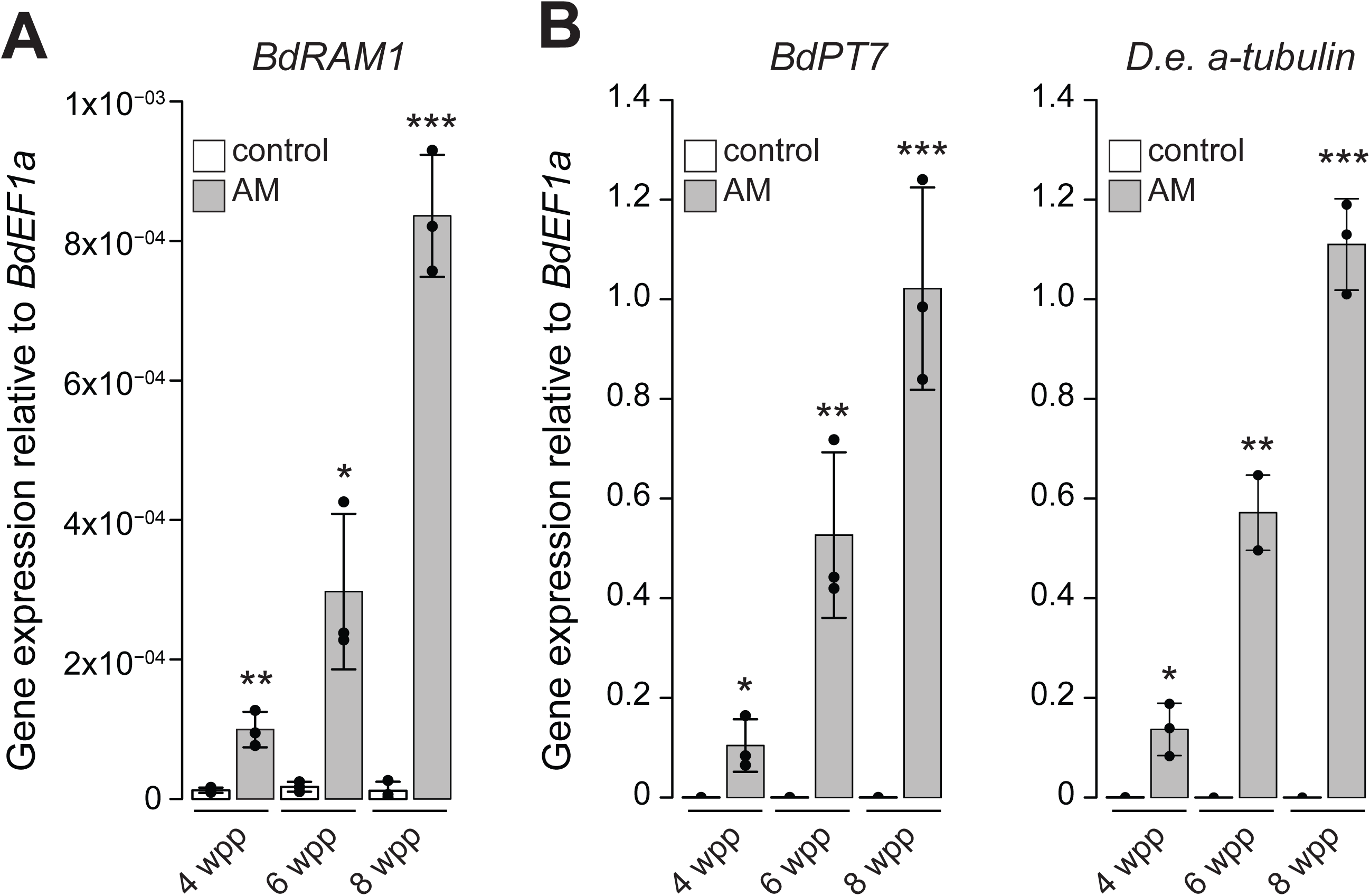
The *B. distachyon* ortholog of *RAM1* is expressed in roots colonized by AM fungi. A) *BdRAM1* gene expression is induced in roots colonized by the AM fungus *D. epigaea* (grey bars, “AM”) relative to non-mycorrhizal, mock-inoculated roots (white bars, “control”). Plants were harvested 4, 6, and 8 weeks post planting (wpp). AM-induced *BdRAM1* gene expression increases over time. B) Gene expression of the AM marker genes *BdPT7* and *D. epigaea (D*.*e*.*) a-tubulin* in *D. epigaea*-colonized and mock-inoculated control roots over time. A) and B) Gene expression was normalized to the *B. distachyon* elongation factor *BdEF1a*. Bar graphs show the mean, error bars the standard deviation. Single points represent individual measurements. Pairwise comparisons of gene expression in AM and control roots were analyzed separately for each time point (Student’s t-test). Significance values: ***p<0.001; **p<0.01; *p<0.05.

Several other GRAS proteins are essential for AM symbiosis including DELLA/SLR1, a negative regulator of GA signaling (Floss et al., 2013; Foo et al., 2013; Yu et al., 2014; Floss et al., 2017). In *della* mutants, AM fungi show a severely reduced ability to enter cortical cells, and as a result almost no arbuscules are formed (Floss et al., 2013; Foo et al., 2013; Yu et al., 2014). Arbuscules are ephemeral structures, and the few arbuscules that are formed in *della* mutants display an increased lifespan, indicating that DELLA not only regulates arbuscule formation but also their degradation (Floss et al., 2017). Two other GRAS transcription factors critical for hormone signaling and AM symbiosis are *NSP1* and *NSP2*. These transcription factors regulate phosphate-dependent strigolactone (SL) biosynthesis in *M. truncatula* and rice (Liu et al., 2011). SLs serve as direct plant communication molecules with AM fungi at the onset of the symbiosis. Mutants impaired in NSP or enzymes required for SL biosynthesis show a reduction in fungal entry into the root and consequently reduced colonization (Gomez-Roldan et al., 2008; Liu et al., 2011; Kobae et al., 2018). Thus, there are several examples of GRAS factors that connect hormone signaling and AM symbiosis.

Many GRAS factors operate in complexes with other GRAS proteins and emerging evidence suggests that this is also true of those involved in AM symbiosis. *M. truncatula* and *L. japonicus* RAM1 were reported to interact with RAD1 and NSP2 (which also interact with each other), but not NSP1 (Gobbato et al., 2012; Park et al., 2015; Xue et al., 2015; Heck et al., 2016). In addition, rice RAM1 interacts with the GRAS transcription factor DIP1, which in turn interacts with DELLA (Yu et al., 2014). *M. truncatula* DELLA proteins were found to interact with numerous other GRAS transcription factors, including RAD1, MIG1, NSP1, and NSP2 (Floss et al., 2016; Fonouni-Farde et al., 2016; Heck et al., 2016; Jin et al., 2016). While their functional significance for symbiosis remains to be determined, the interactions suggest the existence of interconnected transcriptional modules regulated by multiple GRAS transcription factors.

*Brachypodium distachyon* is a monocot model species capable of forming AM symbiosis (Hong et al., 2012) and amenable to genetic manipulation (Bragg et al., 2015). A recent study identified 48 GRAS transcription factors in the genome of *B. distachyon* (Niu et al., 2019). Here we report functional analyses of the GRAS transcription factor *RAM1* in a monocot and assess the potential to alter the levels of symbiotic interfaces by manipulating *RAM1* expression.

## Results and Discussion

We identified *Bradi4g18390* as the single *B. distachyon* homolog of the GRAS transcription factor *RAM1* (Supplemental Fig. 1), a gene that is conserved in AM host plants and missing from non-hosts (Bravo et al., 2016). Similar to orthologous *RAM1* genes of *M. truncatula* (Gobbato et al., 2013; Park et al., 2015), *L. japonicus* (Xue et al., 2015; Pimprikar et al., 2016) and *P. hybrida* (Rich et al., 2015), *B. distachyon RAM1* expression is induced in mycorrhizal roots. Following inoculation with the AM fungus *Diversispora epigaea* (formerly *Glomus versiforme*), *BdRAM1* transcripts increased over time in parallel with increasing colonization of the root system as reported by *D. epigaea a-tubulin* transcripts and the phosphate transporter gene *BdPT7* (Hong et al., 2012), a plant gene marker of AM symbiosis (Figure 1A, B). However, while the transcriptional patterns mirrors the marker genes, it is noticeable that *BdRAM1* transcript levels are low, as is often the case for transcriptional regulators.

The role of *RAM1* in AM has been established in at least three dicot host plants (Gobbato et al., 2013; Park et al., 2015; Rich et al., 2015; Xue et al., 2015; Pimprikar et al., 2016) where it is essential to support arbuscule development and appears to act in the upper tier of a transcription factor hierarchy (Luginbuehl et al., 2017); when ectopically over-expressed in roots, *RAM1* is sufficient to induce expression of several AM-induced genes in the absence of symbiosis (Park et al., 2015; Pimprikar et al., 2016). Given its AM-inducible expression and pivotal regulatory role, we hypothesized that constitutive, high-level expression of *RAM1* might increase the occurrence of arbuscules and possibly overall colonization levels, and this might provide an opportunity to evaluate the functional consequences of modifying colonization patterns. To test this hypothesis, we transformed *B. distachyon* with an overexpression construct, *BdRAM1* under the control of two copies of the constitutively active *CaMV 35S* promoter (*35S:BdRAM1*). In addition, we generated *B. distachyon ram1* loss-of-function mutants via CRISPR/Cas9 editing.

### Arbuscule development in *B. distachyon ram1* mutants is partly impaired

Five independent transgenic lines carrying a CRISPR/CAS9 construct targeting *BdRAM1* were generated and two lines in which *BdRAM1* had been edited were chosen for subsequent analysis. In both transgenic lines, the genome had been edited by both guides (Supplemental Fig. 2); editing by the upstream-most guide resulted in premature stop codons and created a truncated protein of 16 amino acids in the first line, designated *ram1-1*. The second line, designated *ram1-2*, was bi-allelic with edits resulting in premature stop codons that generated truncated protein products of 16 and 42 amino acids.

**Figure 2:**
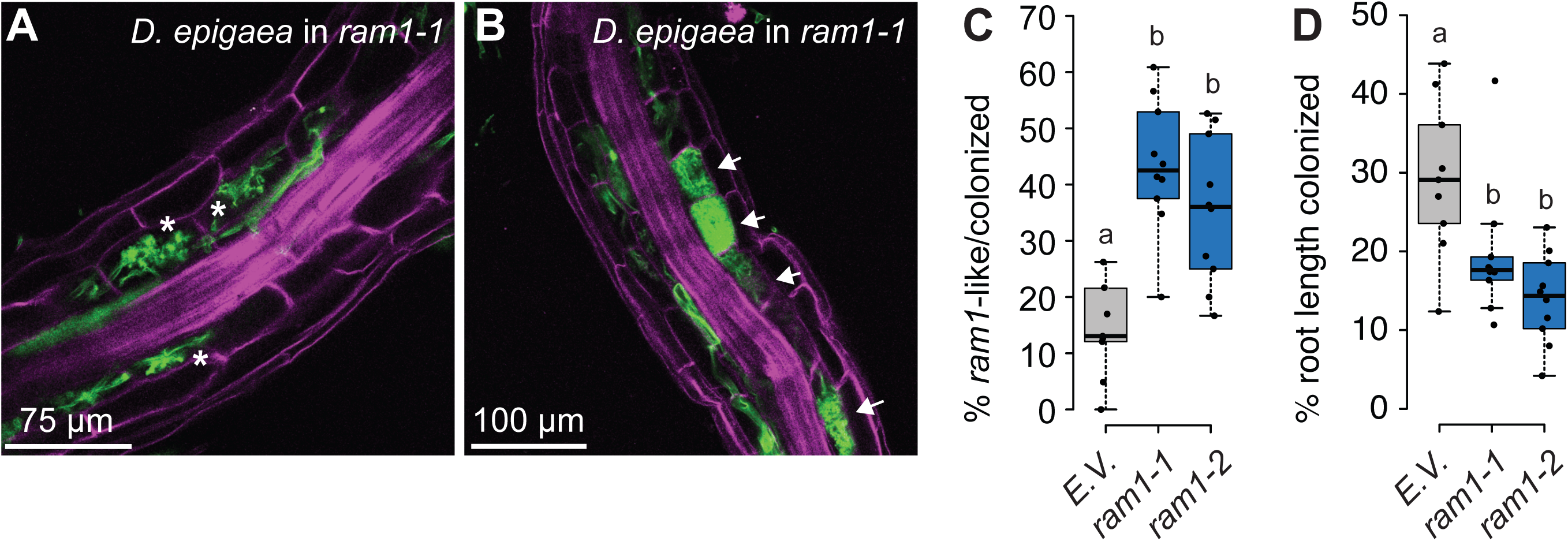
Arbuscule formation is impaired in *B. distachyon ram1* mutants. A) A root piece with “*ram1*-like” *D. epigaea* arbuscules in CRISPR *ram1* mutants. “*Ram1*-like” infections contain solely arbuscules that are not fully developed and show only sparse branching (asterisks). B) A root piece with “wild-type-like” *D. epigaea* arbuscules in CRISPR *ram1* mutants. “Wild-type-like” infections contain fully developed arbuscules (arrows). A) and B), *D. epigaea* fungal structures are visualized using WGA-Alexafluor488 (green), plant cell walls were counter-stained using Propidium Iodide (pink). C) Quantification of “*ram1*-like” infections relative to the total number of infections in *ram1* CRISPR plants and *B. distachyon* plants transformed with the empty vector (E.V.). The proportion of aberrant infections is increased in two *ram1* alleles (ANOVA p=2.38×10^−5^). D) Quantification of total *D. epigaea* root length colonization in CRISPR *ram1* plants relative to E.V.-controls. Root length colonization is significantly decreased in two *ram1* mutant alleles (ANOVA p=0.0014). Pairwise comparisons in C) and D) were performed using Tukey’s HSD post-hoc test; different letters denote significant differences. Box-and-whiskers plots show lower and upper quartiles, and minimum and maximum values. The horizontal bar represents the median, and the points individual measurements. All results presented in this figure were obtained from the T_3_ generation.

*ram1-1* and *ram1-2* were inoculated with *D. epigaea* and the fungal colonization patterns were examined. Some *ram1* roots showed aberrant infection units, reminiscent of the typical dicot *ram1* phenotype, with intraradical hyphae and only small, sparsely branched arbuscules but no fully developed arbuscules (Figure 2A). However, infection units in other roots of the same plant, or even in other parts of the same root, showed an apparent wild-type morphology with some large, well-branched arbuscules (Figure 2B). In comparison with the empty vector control (E.V.) the frequency of aberrant infections in the *B. distachyon ram1* mutants was 2.5 to 3-fold higher and their overall root colonization levels were 34 to 52% lower. Similar results were obtained in several experiments across several generations (Supplemental Fig. 2). These results indicate that *BdRAM1* is required to enable wild-type levels of arbuscule development similar to its orthologs in dicots; however, the *B. distachyon ram1* phenotype is clearly milder than that observed in dicot *ram1* mutants. The finding that *B. distachyon ram1* can support some full arbuscule development suggests that other proteins or pathways have the potential to compensate for loss of *BdRAM1* function. One possible candidate is the GRAS protein *RAD1*, which is closely related to *RAM1* and induced in roots highly colonized by AM fungi (Supplemental Fig. 1, Supplemental Fig. 3). In legumes, there is evidence of a species-specific “micro-diversification” of *RAM1* and *RAD1*, with the relative contributions of the two transcription factors to arbuscule development and symbiotic gene expression varying depending on the host species (Park et al., 2015; Xue et al., 2015; Pimprikar et al., 2016; Pimprikar and Gutjahr, 2018). It is therefore conceivable that some diversification of GRAS factor functions has occurred during the evolution of monocots, which might explain the milder arbuscule development phenotype of *B. distachyon ram1* mutants relative to *M. truncatula ram1*. Interestingly, there are other GRAS factor examples where the converse is true, for example the DELLA proteins, where rice *slr* (Yu et al., 2014) shows a stronger phenotype than the *M. truncatula della* double or triple mutants (Floss et al., 2013; Floss et al., 2017). However, in the absence of other monocot *ram1* mutants for comparison, it is also possible that the milder *ram1* phenotype observed here is a feature specific to *B. distachyon*.

**Figure 3:**
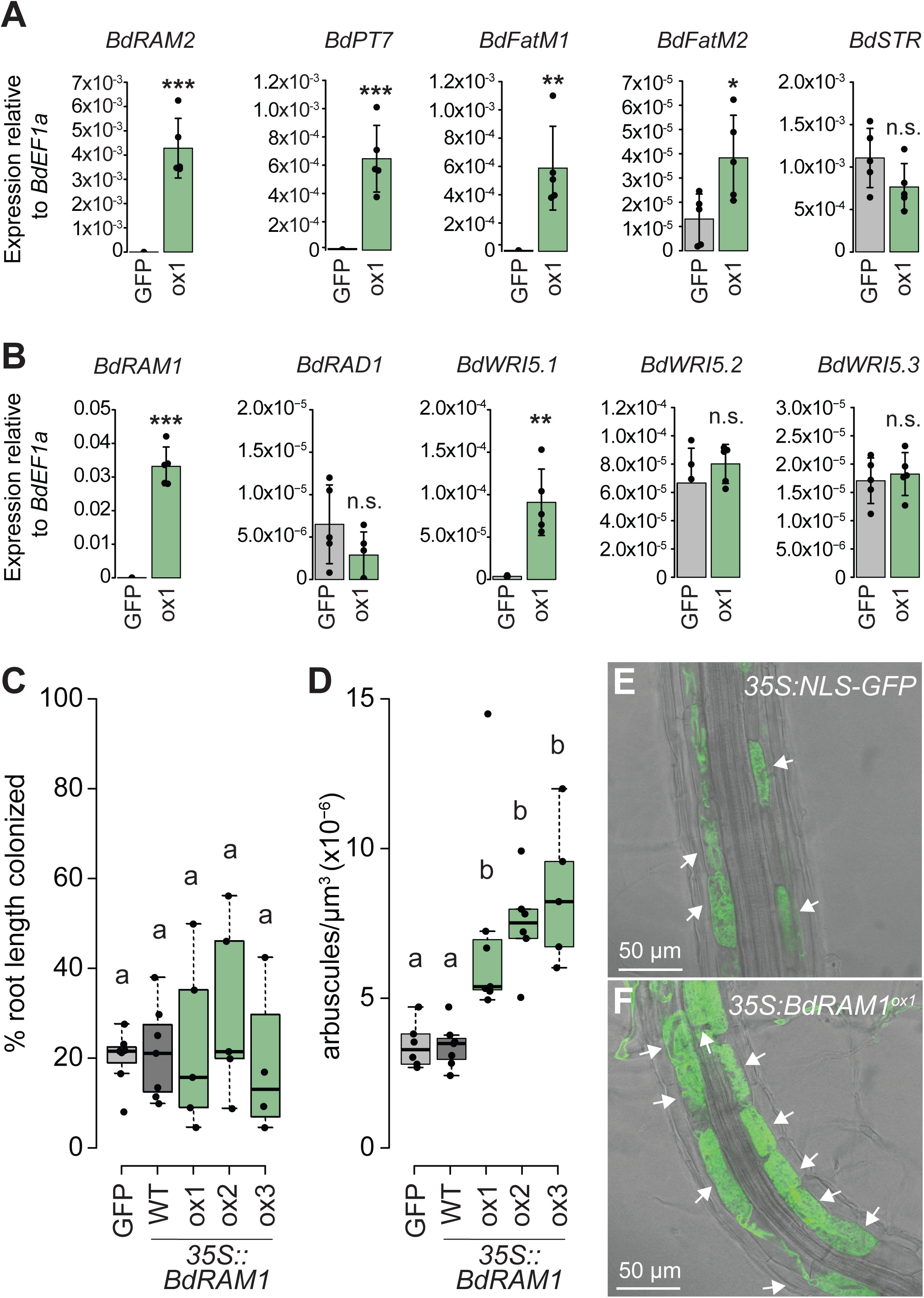
Ectopic overexpression of *BdRAM1* promotes arbuscule formation and expression of AM marker genes. A) Gene expression levels of *B. distachyon* orthologs of *BdRAM1* target genes in non-colonized *35S:NLS-GFP* (denoted as “GFP”) and *35S:BdRAM1*^*ox*^ line #1 (denoted as “ox1”) roots. *35S:BdRAM1*^*ox*^ roots display induced expression of *BdRAM1*, as well as *BdRAM2, BdPT7, BdFatM1*, and *BdFatM2* in the absence of symbiosis relative to *35S:NLS-GFP* control roots. *BdSTR* gene expression is not affected in these roots. B) Gene expression of *B. distachyon RAD1* and *WRI5* orthologs. Only expression of *BdWRI5*.*1* is induced in non-colonized roots over-expressing *BdRAM1* (*35S:BdRAM1*^*ox*^) relative to *35S:NLS-GFP* control roots. A), B), Bar graphs show the mean, error bars the standard deviation. Single points represent individual measurements. Pairwise comparisons were estimated using the Student’s t-test. Significance codes: ***p<0.001; **p<0.01; *p<0.05; n.s., not significant. C) Quantification of total root colonization in independent lines transformed with *35S:BdRAM1*. There is no difference in overall root colonization between three lines ectopically over-expressing *35S:BdRAM1* (*35S:BdRAM1*^*ox*^, denoted as “ox1”, “ox2”, “ox3”) and control plants (*35S:BdRAM1*^*WT*^, which does not over-express *BdRAM1* and was therefore denoted as “WT”; and *35S:NLS-GFP*, labeled as “GFP”). ANOVA p=0.71. Root length colonization was quantified using the grid-line method (McGonigle et al., 1990). D) Quantification of *D. epigaea* arbuscules in a defined area at the fungal appressorium. Roots of 3 independent transgenic *35S:BdRAM1*^*ox*^ lines (denoted as “ox1”, “ox2”, ox3”) contain more arbuscules than roots transformed with the control construct *35S:NLS-GFP* (“GFP”), or roots that contain *35S:BdRAM1* but do not overexpress the gene (“WT”). Arbuscule number was normalized to the volume of the confocal stack. Kruskal-Wallis test p=1.32×10^−4^, pairwise comparisons were conducted using the Dunn’s post-hoc test; different letters denote significant differences. C) and D), Box-and-whiskers plots show lower and upper quartiles, and minimum and maximum values. The horizontal bar represents the median, and the points individual measurements. E) Representative image of a *35S:NLS-GFP* root colonized by *D. epigaea*. F) Representative image of a *RAM1*-overexpressing *35S:BdRAM1*^*ox*^ (line #1) root colonized by *D. epigaea*. E) and F), arbuscules are highlighted with arrows.

### Overexpression of *RAM1* alters plant morphology and results in constitutive expression of AM marker genes

The generation of transgenic *B. distachyon* plants overexpressing *RAM1* was surprisingly challenging; from two, full-scale independent transformation experiments, only three viable independent transgenic *35S*:*BdRAM1* overexpressing lines (*35S:BdRAM1*^*ox*^) were obtained and the seed production from these lines was exceedingly poor. In addition, we obtained two lines, which carried the *35S:BdRAM1 T-DNA* but displayed wild type-like *BdRAM1* transcript levels (*35S:BdRAM1*^*WT*^). By contrast, transgenic plants carrying *35S:NLS-GFP-GUS* (hereto referred to simply as *35S:NLS-GFP)*, were generated without difficulty. Seed production from the latter two genotypes was not impaired.

In addition to poor seed production and viability, the shoot and root phenotypes of the three lines with transcriptional up-regulation of *BdRAM1* (*35S:BdRAM1*^*ox*^) differed from the vector controls and from the *35S:BdRAM1*^*WT*^ plants. The *35S:BdRAM1*^*ox*^ plants were characterized by a stunted bushy shoot with increased tiller formation and increased leaf angles, as well as a decreased number of node roots (Figure 4A, Supplemental Fig. 4). Thus, constitutive, ectopic, overexpression of *BdRAM1*, clearly influences plant development. While the cause is unknown, it might be the result of ectopic expression of *BdRAM1* target genes and/or perhaps an interference of RAM1 with other GRAS transcriptional networks, many of which regulate development (reviewed in (Cenci and Rouard, 2017). Either way, the aberrant developmental phenotypes likely explain the difficulties in regenerating transgenic lines and their fecundity.

**Figure 4:**
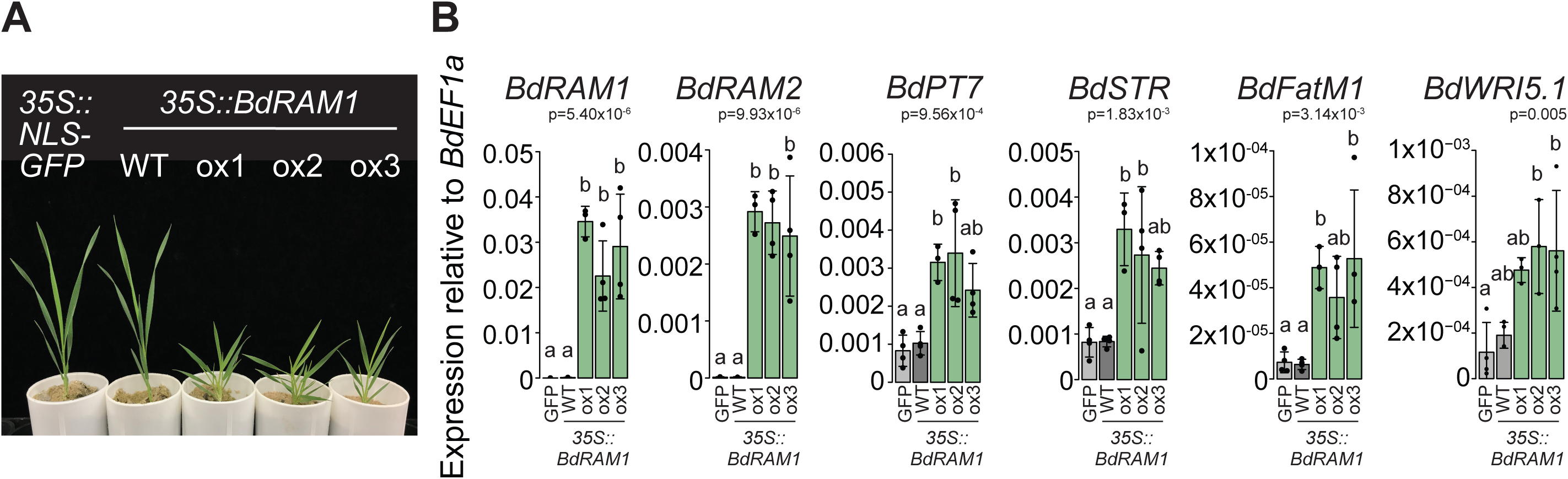
*RAM1* overexpressors show altered shoot development and constitutively express root AM marker genes in their shoots. A) Photograph of 4.5 week-old *B. distachyon* plants transformed with *35S:NLS-GFP* or *35S:BdRAM1*. The three independent transformant lines overexpressing *BdRAM1* (“ox1”, “ox2”, “ox3”) display a bushy stature, whereas the *35S:BdRAM1*-transformant line not overexpressing *BdRAM1* (“WT”) resembles the *35S:NLS-GFP* control plant. B) Gene expression of *BdRAM1* and of several root AM marker genes and *BdWRI5*.*1* is strongly induced in in 4.5-week shoots of three *35S:BdRAM1*^*ox*^ lines (“ox1”, “ox2”, “ox3”) relative to control plants transformed with *35S:NLS-GFP* (“GFP”) or the *35S:BdRAM1*-transformant line not overexpressing *BdRAM1* (“WT”). Bar graphs show the mean, error bars the standard deviation. Single points represent individual measurements. Significance values (ANOVA) for each gene are indicated in the figure. Pairwise comparisons were conducted using Tukey’s HSD post-hoc test; different letters denote significant differences.

The *35S:BdRAM1*^*ox*^ plants showed constitutive expression of *B. distachyon* orthologs of *RAM2, STR, PT4 and FatM* (Figure 3A); elevated expression of these genes would normally occur only in response to colonization by AMF (for example, Harrison et al., 2002; Paszkowski et al., 2002; Gutjahr et al., 2008; Zhang et al., 2010; Gobbato et al., 2012; Gutjahr et al., 2012; Hong et al., 2012; Bravo et al., 2017), and we observed a similar expression pattern of their orthologs in *B. distachyon* roots (Figure 1B, Supplementary Fig. 3). Given the prior knowledge from dicots, we had anticipated that *35S:BdRAM1*^*ox*^ would increase expression of these genes in roots, but it was surprising to see that expression of these genes was also induced in shoots (Figure 4B). There were some exceptions, expression of *BdSTR* increased in *35S:BdRAM1*^*ox*^ shoots, but not in roots while *BdFatM2* showed the opposite expression pattern (Figure 3A, Figure 4B, Supplemental Fig. 5). Overall, these data indicate that *BdRAM1* alone is sufficient to drive increased expression of these genes in the absence of AM fungi and that transcription co-factors - if required for *Bd*RAM1 function - must be present in all tissues.

**Figure 5:**
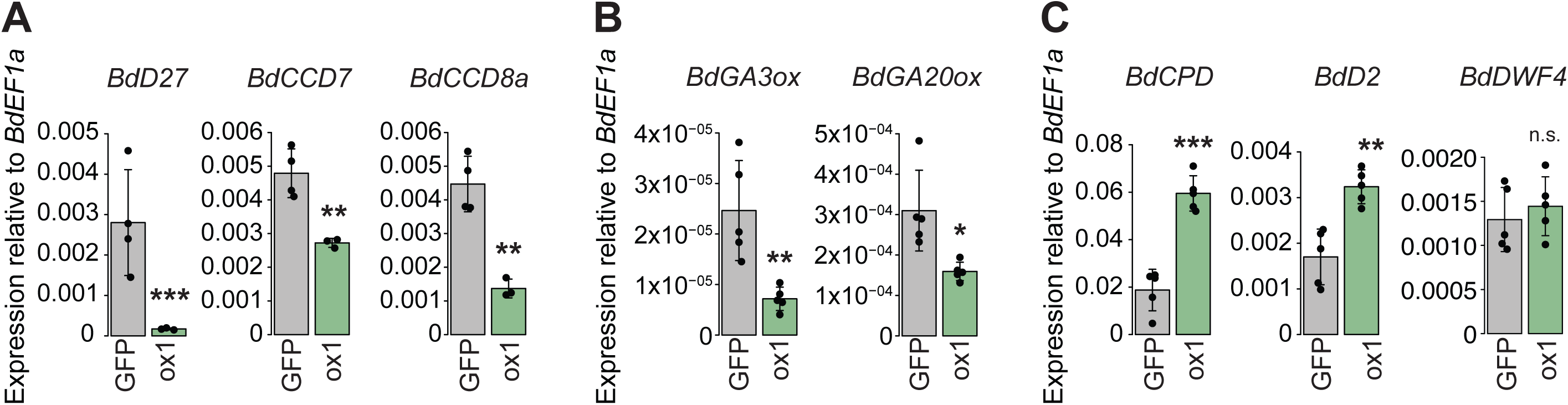
Expression of hormone biosynthesis genes is altered in the *RAM1* overexpressors. A) *B. distachyon* orthologs of three genes involved in the Strigolactone biosynthesis pathway (*BdD27, BdD17, Bd10*) are down-regulated in non-colonized roots ectopically overexpressing *BdRAM1* (*35S:BdRAM1*^*ox*^ line#1, denoted as “ox1”) relative to *35S:NLS-GFP* (“GFP”) control roots. B) Two genes with a putative function in Gibberellic acid biosynthesis (*BdGA3ox1, BdGA20ox1*) are down-regulated in *35S:BdRAM1*^*ox*^ roots. C) Two *B. distachyon* genes orthologous to known Brassinosteroid biosynthesis genes (*BdCPD, BdD2/BdCYP91D*) are induced in *35S:BdRAM1*^*ox*^ roots. A third gene, *BdDWF4*, is not affected. Bar graphs show the mean, error bars the standard deviation. Single points represent individual measurements. Pairwise comparisons were estimated using the Student’s t-test. Significance codes: ***p<0.001; **p<0.01; *p<0.05; n.s., not significant.

In dicots, *RAM1* regulates expression of a second tier of transcription factors including *RAD1* and three members of the *WRINKLED* family (*WRI5a-c*); the latter directly regulate expression of lipid genes (Park et al., 2015; Jiang et al., 2017; Luginbuehl et al., 2017). We found that *BdRAD1*, and the three *B. distachyon* AP2 family transcription factors most closely related to *MtWRI5a-c* (further denoted as *BdWRI5*.*1, BdWRI5*.*2, and BdWRI5*.*3*) were strongly induced in wild-type roots colonized with *D. epigaea* relative to mock-inoculated controls (Supplemental Fig. 3) but interestingly, only *BdWRI5*.*1* was induced in non-colonized *35S:BdRAM1*^*ox*^ roots (Figure 3B). A similar pattern was observed in *35S:BdRAM1*^*ox*^ shoots (Figure 4B, Supplemental Fig. 5). Thus, in contrast to *M. truncatula*, a RAM1-independent pathway likely leads to up-regulation of *BdRAD1, BdWRI5*.*2*, and *BdWRI5*.*3* in mycorrhizal roots. This points to functional diversification of the regulatory cascade responsible for the transcriptional reprogramming of roots during AM symbiosis in *B. distachyon*. In addition, it may provide an explanation for the relatively mild *ram1* mutant phenotype we observed (Figure 2). Future research in other monocot species is required to determine if such a functional diversification is unique to *B. distachyon* or a monocot-specific phenomenon.

### Arbuscule density is higher in *RAM1* overexpressors relative to controls

The initial goal of this study was to test the hypothesis that constitutive overexpression of *RAM1* would increase arbuscule density and/or colonization and then to use the plants to address secondary hypotheses about symbiotic performance.

To test the first hypothesis, we grew *35S:BdRAM1*^*ox*^, *35S:BdRAM1*^*WT*^ and *35S:NLS-GFP* control plants in substrate containing *D. epigaea* spores and evaluated colonization levels and arbuscule morphology. Colonization levels in *35S:BdRAM1*^*ox*^ and control plants did not differ significantly, although the variation was much greater in the *35S:BdRAM1*^*ox*^ plants (Figure 3C). Arbuscules in *35S:BdRAM1*^*ox*^ plants showed a wild-type morphology, but the number of arbuscules, which we assessed within a defined root volume below the hyphopodium, was on average 2-fold greater in the *35S:BdRAM*^*ox*^ plants relative to controls (Figure 3D-F, Supplemental Fig. 5). Thus, *35S:BdRAM1*^*ox*^ plants have a higher capacity to establish and/or to maintain arbuscules relative to the control plants. As RAM1 regulates the expression of several other transcription factors, as well as genes involved in lipid biosynthesis and nutrient transport, the increased arbuscule density in the *35S:BdRAM1*^*ox*^ plants may result from a combination of factors including arbuscule initiation and/or regulation of arbuscule lifespan.

Unfortunately, the severe shoot growth and branching phenotype of the *BdRAM1* overexpressors prevented a fair evaluation of symbiotic performance (Supplemental Fig. 5). While colonized *35S:BdRAM1*^*ox*^ plants and controls both showed an increase in shoot fresh weight and tiller number relative to their respective mock-inoculated controls, the differences in the developmental architecture of these lines precluded direct physiological comparisons. Consequently, it was not possible to determine whether the increased arbuscule density influenced symbiotic performance.

### Hormone biosynthetic and regulatory gene expression is altered in *BdRAM1* overexpressors

The shoot architecture phenotype of the *35S:BdRAM1*^*ox*^ plants is reminiscent of the phenotypes of several monocot hormone mutants. For example, rice and *B. distachyon* mutants defective in GA, SL, and BR biosynthesis or signaling display dwarf phenotypes with increased tillering (e.g. (Spielmeyer et al., 2002; Ishikawa et al., 2005; Asano et al., 2009; Lin et al., 2009; Thole et al., 2012). To obtain further clues about the *BdRAM1* overexpression phenotype, we evaluated the expression of several genes associated with SL, GA, and BR signaling. *B. distachyon* orthologues of genes involved in SL biosynthesis (*BdD27, BdD17, BdD10*) (Seto and Yamaguchi, 2014) and GA biosynthesis (potential orthologs of *A. thaliana GA3ox1* and *GA20ox1*) (Kakei et al., 2015) were down-regulated, while key BR biosynthesis genes (*BdCPD, BdD2/CYP90D*) but not *BdDWF4* (Kakei et al., 2015) were elevated in non-colonized *35S:BdRAM1*^*ox*^ roots relative to the controls (Figure 5A, B). In contrast, the GA receptor *GID1* and the GA-regulator *DELLA/SLR1* (Daviere and Achard, 2013) as well as the regulators of SL signaling *D3* and *D53* (Seto and Yamaguchi, 2014), and the *B. distachyon* BR receptor *BdBRI1* and the BR-responsive transcription factor *BdBZR1* (Corvalan and Choe, 2017) were differentially regulated in *35S:BdRAM1*^*ox*^ roots relative to controls (Supplemental Fig. 6). Thus the transcript data indicate a disturbance in hormone biosynthetic and regulatory gene expression likely contributing to the altered shoot architecture. Because of substantial cross-talk between hormone signaling pathways (Itoh et al., 2001; Umehara et al., 2008; Unterholzner et al., 2015; Corvalan and Choe, 2017), it is not possible to predict the initial cause. As GA, SL and BR hormone pathways each involve regulation via GRAS-transcription factors (Tong et al., 2009; Liu et al., 2011; Chen et al., 2013), it is possible that ectopic overexpression of *BdRAM1* disturbs GRAS-factor complexes, leading to mis-regulation of these pathways. Alternatively, one of the native functions of *BdRAM1* may be to regulate aspects of hormone signaling. For example, in rice, RAM1 interacts with a DELLA interacting protein, DIP, and therefore it is possible that one of RAM1’s native functions is to influence GA signaling and that this is exacerbated in the *35S:BdRAM1*^*ox*^, leading to further downstream effects on other pathways. If mis-regulation of GA biosynthesis gene expression translates to disturbed GA homeostasis in *35S:BdRAM1*^*ox*^ roots, an imbalance in GA-regulated arbuscule formation and degradation could result (Floss et al., 2013; Floss et al., 2017). Such a scenario might also explain the increased arbuscule numbers in *35S:BdRAM1*^*ox*^ roots as well as a dwarf shoot phenotype.

### Conclusion

In conclusion, *BdRAM1*, similar to its orthologs in dicots, regulates arbuscule development and transcriptional regulation of several AM symbiosis-induced genes, although it is likely that there is some functional redundancy with other GRAS or WRI5 transcription factors. Constitutive overexpression of *35S:BdRAM1* increased the number of arbuscules relative to control plants; although the plants were unsuitable for experiments to assess the functional consequences of increasing the symbiotic interfaces, the data nevertheless indicate that it is possible to manipulate arbuscule density through expression of *RAM1*. Future research should focus on increasing *RAM1* gene expression specifically in the root cortex. We predict such a strategy would increase arbuscule numbers without the accompanying developmental defects and would enable evaluation of the consequences of increasing the density of symbiotic interfaces and the effects on nutrient exchange during AM symbiosis.

## Materials and Methods

### Plant material and growth conditions

*B. distachyon* plants were grown in a growth chamber under a 12 h light (24°C)/12 h dark (22°C) regime. For all experiments that were conducted in the absence of an AM fungal symbiont, *B. distachyon* plants were grown in 20.5 cm long cones filled with sterile Terragreen (Oli-Dri) and play sand (Quikrete) in a ratio of 1:1. For all experiments involving AM symbiosis, *B. distachyon* plants were grown in cones filled with a sand-gravel mix, and were inoculated with 250 *Diversispora epigaea* spores (formerly *Glomus versiforme*) as previously described (Muller et al., 2019). For mock-inoculated controls, we added an appropriate volume of filtered spore wash solution instead of the spores. Unless otherwise stated, *B. distachyon* plants were fertilized once per week with 1/4-strength Hoagland’s fertilizer containing 20μM Pi and harvested 4-5 weeks after transplanting to cones.

To monitor AM growth responses, seedlings were planted into pots (3 seedling per 11cm diameter pot and 8 pots per genotype) containing a 1:20 mixture of autoclaved N7/N8 soil (Watts-Williams et al., 2019) to sand/gravel mix. The sand/gravel mix is a 2:2:1 mixture of play sand, fine black sand and gravel (as described in (Floss et al., 2017). 250 surface sterilized *D. epigaea* spores were place below each plant. Beginnning at 3 weeks post planting, the pots were fertilized weekly with 50 ml of 1/4-strength Hoaglands solution lacking phosphate and 9 ml of 0.5mM Ca_3_(PO_4_)_2._ Plants were harvested at 9 weeks post planting. The growth chamber conditions were as described above.

### Plasmid generation

To clone the CRISPR/Cas9 construct targeting *BdRAM1*, we used the vector and cloning system described by Xie et al. (Xie et al., 2015). To design the primers (shown in Supplemental Table 1), gene-specific guide RNA sequences targeting *Bradi4g18390* were identified using CRISPR-P (Lei et al., 2014) and CRISPR-PLANT (Xie et al., 2015) and selected based on their location in the coding sequence and low number of off-target sites. We generated a 2-guide CRISPR/Cas9 construct that targeted *Bradi4g18390* at positions 32-54 bp (guide RNA1) and 280-302 bp (guide RNA2) downstream of the transcription start site (Supplemental Fig. 2). As a negative control we used the empty vector *pRGEB32* (Xie et al., 2015).

To clone *35S:BdRAM1* overexpression constructs, the coding sequence of *Bradi4g18390* was amplified using gene-specific primers flanked by *att*B1 and *att*B2 recombination sites (Supplemental Table 1), and cloned into *pDONR221*, resulting in the *pENTR1-2 BdRAM1* entry clone. *pENTR1-2* clones containing the coding sequence of *NLS-GFP-GUS*, as well as *pENTR4-1* entry clones containing the *CaMV35S* promoter and *pENTR2-3* containing the *CaMV35S* terminator were cloned previously (Ivanov and Harrison, 2014; Floss et al., 2017). To assemble the binary vectors for *B. distachyon* transformation, four vectors (*pENTR4-1* containing the double *CaMV35S* promoter, *pENTR1-2* containing *BdRAM1* or *NLS-GFP-GUS, pENTR2-3* containing the *CaMV35S* terminator and *pHb7m34GW* (Karimi et al., 2005)), were combined to generate *35S:BdRAM1* or *35S:NLS-GFP* using the multi-site gateway cloning system (Invitrogen). All vector sequences were confirmed by Sanger sequencing.

### Generation of *B. distachyon* transformants

The CRISPR/Cas9 constructs targeting *BdRAM1* as well as the *35S:BdRAM1* and *35S:NLS-GFP* constructs were transformed into *B. distachyon* (accession Bd21-3) following a previously established protocol(Bragg et al., 2015). Plantlets emerging from transformed calli (selectable marker: Hygromycin) were transplanted into Metro-Mix 350 and genotyped to test for the presence of the construct (see Supplemental table 1 for primer sequences). In addition, in the case of the CRISPR/Cas9 constructs, the CRISPR/Cas9 target loci were amplified using flanking primers and purified PCR products were Sanger-sequenced in order to identify gene edits.

### Visualization and quantification of fungal root colonization

Fungal colonization of *B. distachyon* roots was visualized by staining with Wheat-Germ Agglutinin (WGA) coupled to Alexafluor488 as previously described (Hong et al., 2012). Roots were observed using a Leica M205 stereomicroscope and root colonization was quantified using the gridline-intersect method (McGonigle et al., 1990). To quantify the *ram1* phenotype, roots intersecting the gridlines were scored into one of three categories: (1), not colonized; (2), colonized with wildtype-like arbuscules; (3), colonized with aberrant (sparsely branched or collapsed arbuscules) or no arbuscules. The ratio of category 3 over the overall number of intersections of colonized roots (category 2+3) x100 was used to determine the percentage of intersections without arbuscules/total colonization. Total root length colonization was calculated as the percentage of category 2+3 over the total number of intersections counted x 100. To study arbuscule morphology, WGA-Alexafluor488-stained roots were counterstained with propidium iodide to visualize plant cell walls, and observed with a Leica SP5 confocal microscope. To quantify arbuscule numbers in *35S:BdRAM1*^*ox*^ roots, confocal stacks from highly colonized roots were taken so that the fungal hyphopodium was in the center of the image to ensure we capture infections of similar developmental stages. The total number of arbuscules per stack was assessed manually using the Fiji Image Analysis Package (Schindelin et al., 2012). Stack depth (z-plane) was chosen to encompass the whole infection, and arbuscule numbers were normalized against the stack volume (length of x*y*z planes). To avoid potentially confounding effects caused by different *B. distachyon* root types, we selected only thin lateral roots with a single layer of cortical cells for analysis.

### RNA isolation, cDNA synthesis, and quantitative PCR

RNA isolation, cDNA synthesis, and quantitative PCR were performed as previously described (Muller et al., 2019). Primers used to quantify expression of target genes are shown in Supplemental table 1. Ct values of the tested genes were normalized against *BdEF1a* {resulting in Ct} and relative expression levels were calculated with the formula 2^-Ct^.

### Assessment of plant morphology

Plants were grown in the absence of AM fungi and whole plants were harvested 2, 4, and 6 weeks after planting. Tiller and node root numbers were counted, and maximal root system and shoot length were measured. The angle between individual leaves and the stem was measured on images of the same plants using the Fiji Image Analysis Package (Schindelin et al., 2012).

### Phylogenetic analyses

*B. distachyon* orthologs of *M. truncatula RAM1, RAD1, RAM2, PT4, FatM, STR, WRI5a-c* as well as *O. sativa D27, D17/CCD7, CCD8a, D3, D53, D14, D14L and SLR1* were identified using phylogenetic approaches described before (Supplemental Fig. 1, 7)(Bravo et al., 2016) *B. distachyon* genes putatively involved in Brassinosteroid and Gibberellic acid biosynthesis and signaling were identified previously (Kakei et al., 2015; Corvalan and Choe, 2017; Niu et al., 2019).

### Statistical analyses and data representation

All experiments were performed using biological replicates. All experiments were repeated at least two times. The distribution of residuals was tested for normality using the Shapiro-Wilk test. If normality assumption was met, pairwise comparisons were analyzed using a two-sided Student’s t-test. For multiple comparisons, the raw data was subjected to a one-way analysis of variance (ANOVA) followed by Tukey’s post-hoc test. If normality assumption was not met, data were analyzed using the Kruskal-Wallis test followed by Dunn’s post-hoc test (p-values adjusted after Benjamini-Hochberg). All statistical analyses were performed using R software. Quantification data for n>5 biological replicates are represented as box-and-whiskers plots, which show the lower and upper quartiles as well as the minimum and maximum values. The horizontal line in the box plots represents the median. Points represent single measurements. For datasets with less than 5 measurements per genotype, bar plots were chosen. Bars represent the mean, and error bars the standard deviation. Points represent single measurements.

## Acknowledgements

We thank Sophia Cotraccia, Cassandra Proctor, and Stephanie Roh for technical assistance and the BTI Biotechnology center for generating some of the *B. distachyon* transgenic lines. Financial support for the project was provided by the U.S. Department of Energy, Office of Science, Office of Biological and Environmental Research (grant no. DE-SC0012460) and the TRIAD Foundation. LMM was supported by Postdoctoral Fellowships from the Swiss National Science Foundation (SNF, Early Postdoc.Mobility) and the German Research Foundation (DFG), LCS was supported by a Marie Curie Fellowship (FP7-PEOPLE-2013-IOF-624739).

## Figure legends

**Figure S1:**
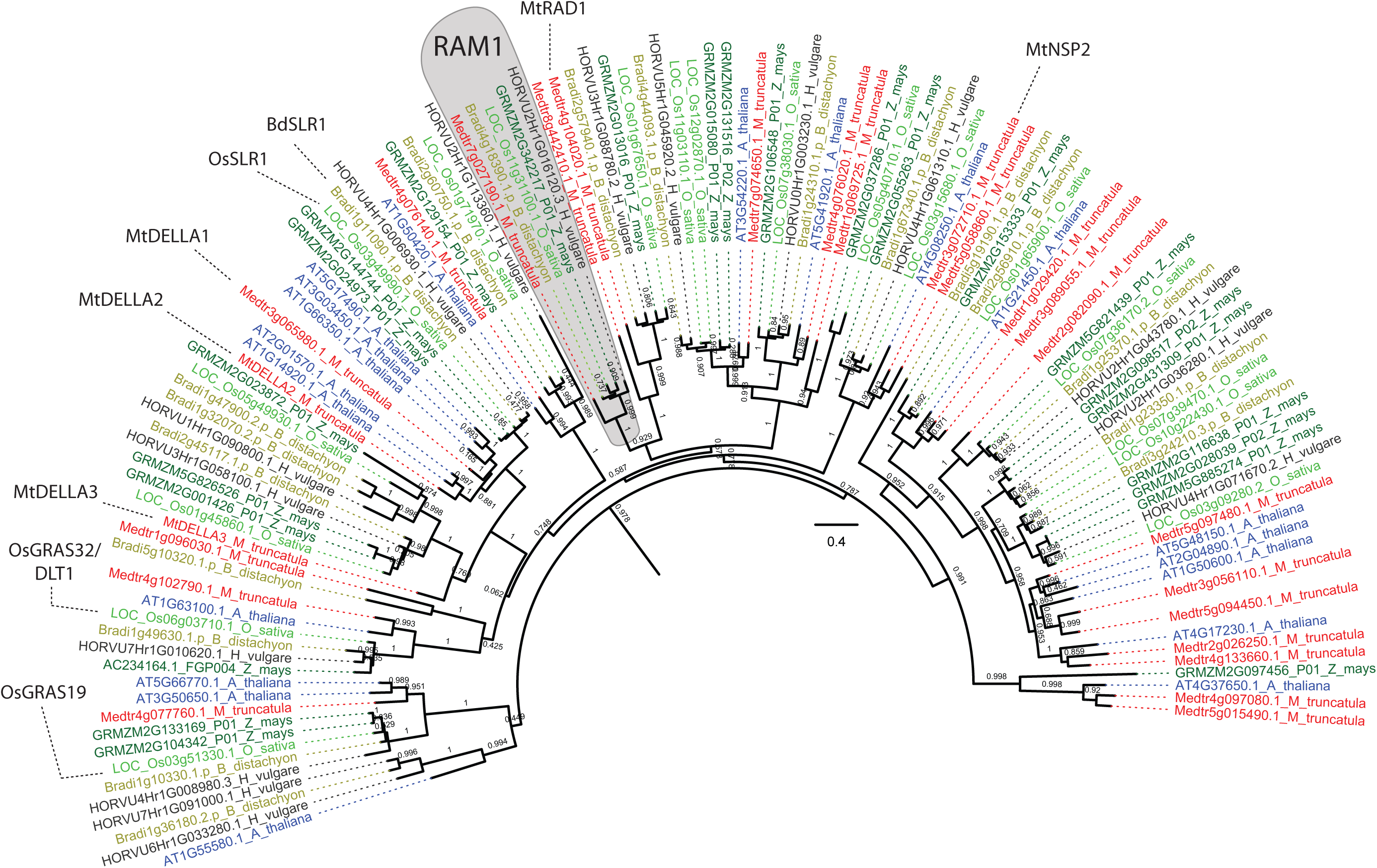
Phylogenetic tree showing GRAS transcription factors related to *Bd*RAM1 in the AM host species (*B. distachyon, O. sativa, H. vulgare, Z. mays, M. truncatula*) and the non-host species *A. thaliana*. The phylogeny is based on the *Bd*RAM1 protein sequence and was constructed using a previously published pipeline(Bravo et al., 2016). *Mt*RAD1 regulates arbuscule formation(Park et al., 2015). *Os*GRAS19 and *Os*GRAS32 function in Brassinosteroid signaling (Tong et al., 2009; Chen et al., 2013; Floss et al., 2013; Floss et al., 2016) and *Mt*DELLA function in Gibberellic acid signaling and AM symbiosis (Floss et al., 2013; Yu et al., 2014; Floss et al., 2017). *Mt*NSP2 regulates strigolactone biosynthesis and AM symbiosis (Liu et al., 2011).

**Figure S2:**
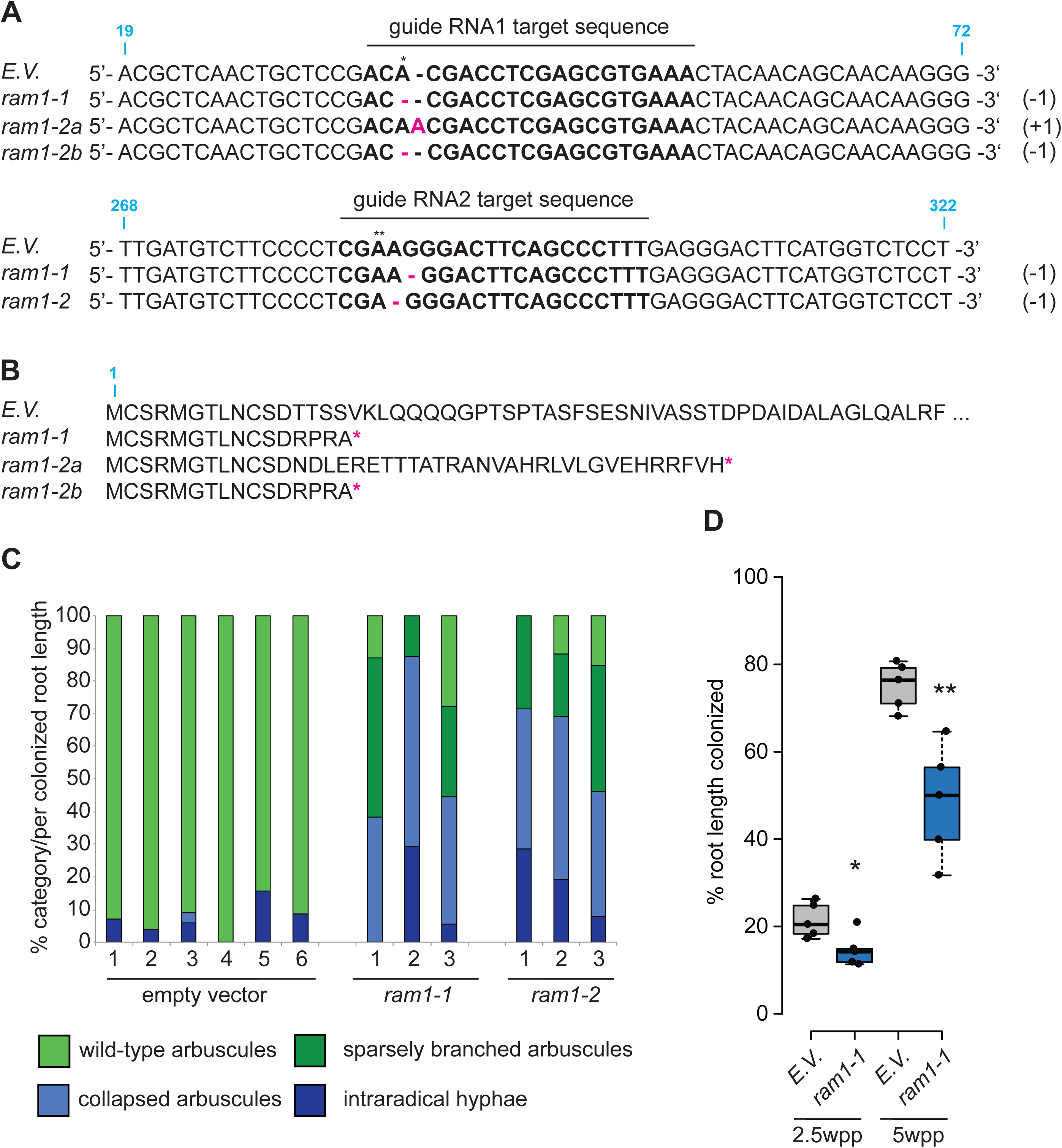
*B. distachyon* CRISPR/Cas9 edited *ram1* mutants. A) Alignments of *Bdram1* DNA sequences in *ram1-1* and *ram1-2* lines, as well as in *B. distachyon* plants transformed with the empty-vector control (E.V.). Target sequences of the two guide RNAs are indicated, as well as CRISPR/Cas9-induced insertions and deletions (pink, T_3_ generation). Numbers in blue indicate nucleotide position in the wild-type *Bdram1* coding sequence. We detected a homozygous single-base pair deletion in *ram1-1* at the target sequence of the upstream-most guide. In *ram1-2*, two edited variants (a one base-pair insertion in one allele, and a one base-pair deletion in the other; denoted as a and b, respectively) were detected at the target sequence of guideRNAl. Both *ram1-1* and *ram1-2* showed homozygous edits caused by guideRNA2 further downstream in the gene. B) All indels caused by guideRNAl lead to a frameshift and a premature stop codon (pink asterisk) after amino acid positions l6 or 42, respectively. C) Morphologic analysis of root colonization phenotypes in T_l_ *ram1-1* and *ram1-2* plants compared to empty vector controls. Both *ram1* alleles display increased numbers of sparsely branched or collapsed arbuscules and infections without any arbuscules relative to control plants, where most arbuscules were of wildtype-like morphology. Each bar represents an individual root system; fungal colonization patterns were divided in four categories (color-coded) and counted microscopically using the grid-line method (McGonigle et al., l990). No difference in overall root colonization was observed in this experiment. D) Overall root colonization of *B. distachyon ram1-1* mutants (T_2_ generation) is reduced relative to empty-vector controls (E.V.). Plants were harvested at two time points: 2.5 and 5 weeks post planting (wpp). Pairwise comparisons were calculated separately for each time point (Student’s t-test). Significance codes: **p<0.0l; *p<0.05. Arbuscule phenotypes were not analyzed in this experiment.

**Figure S3:**
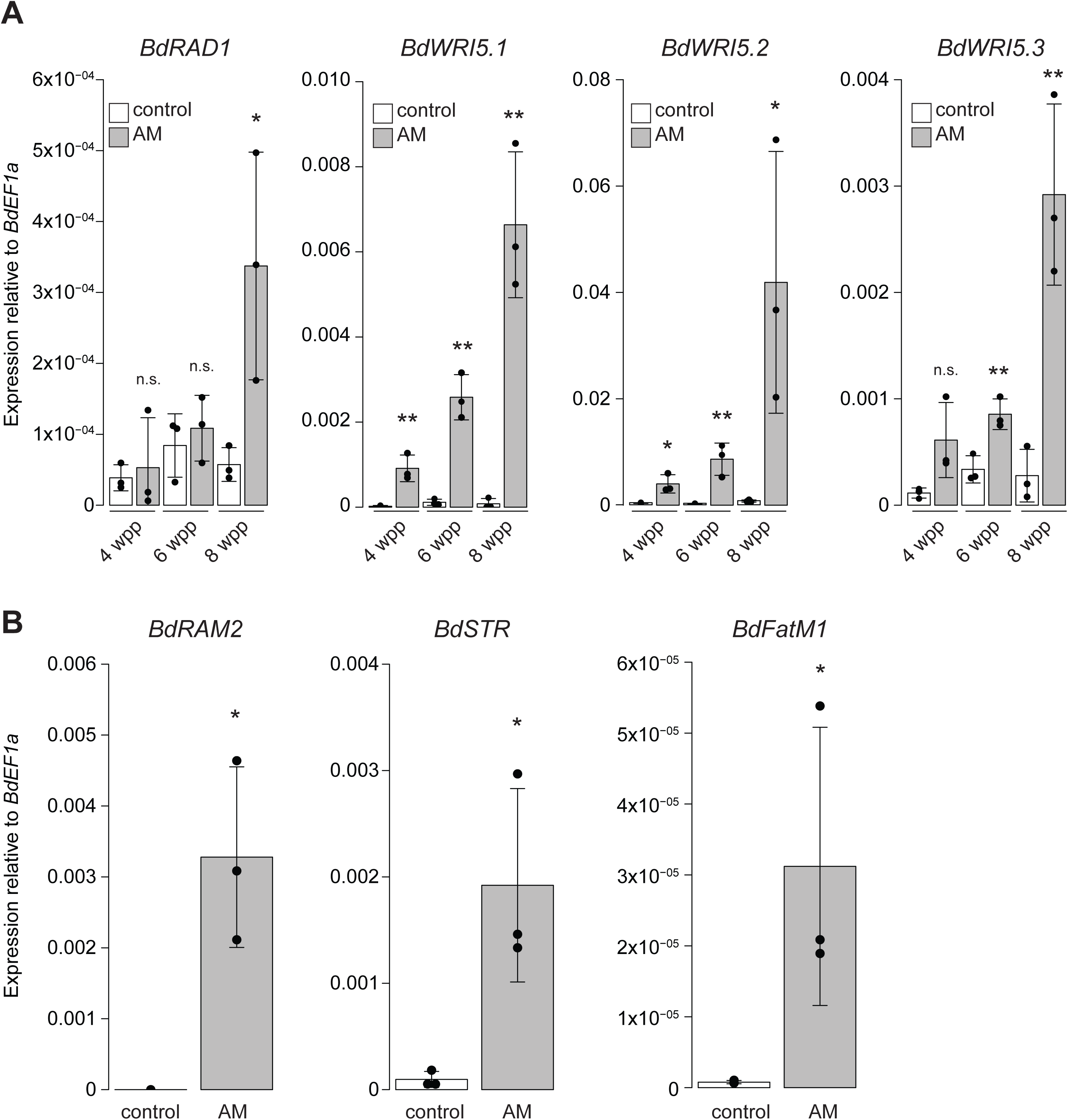
Gene expression of selected genes in colonized roots. A) Gene expression of the GRAS transcription factor *BdRADl* and three AP2-family transcription factors related to *MtWR/5*. Gene expression was measured in mock-inoculated roots (“control”) and roots colonized with *D. epigaea* (“AM”) harvested at 4, 6, and 8 weeks after planting (wpp). Pairwise comparisons between AM and control roots were calculated separately for each time point (Student’s t-test). B) Gene expression of selected *B. distachyon* orthologs of the AM marker genes *RAM2, STR*, and *FatM* in 6 week-old *D. epigaea*-colonized (“AM”) roots relative to mock-inoculated control roots. Pairwise differences were calculated using Student’s t-test. Significance codes in A) and B): ***p<0.00l; **p<0.0l; *p<0.05; n.s., not significant.

**Figure S4:**
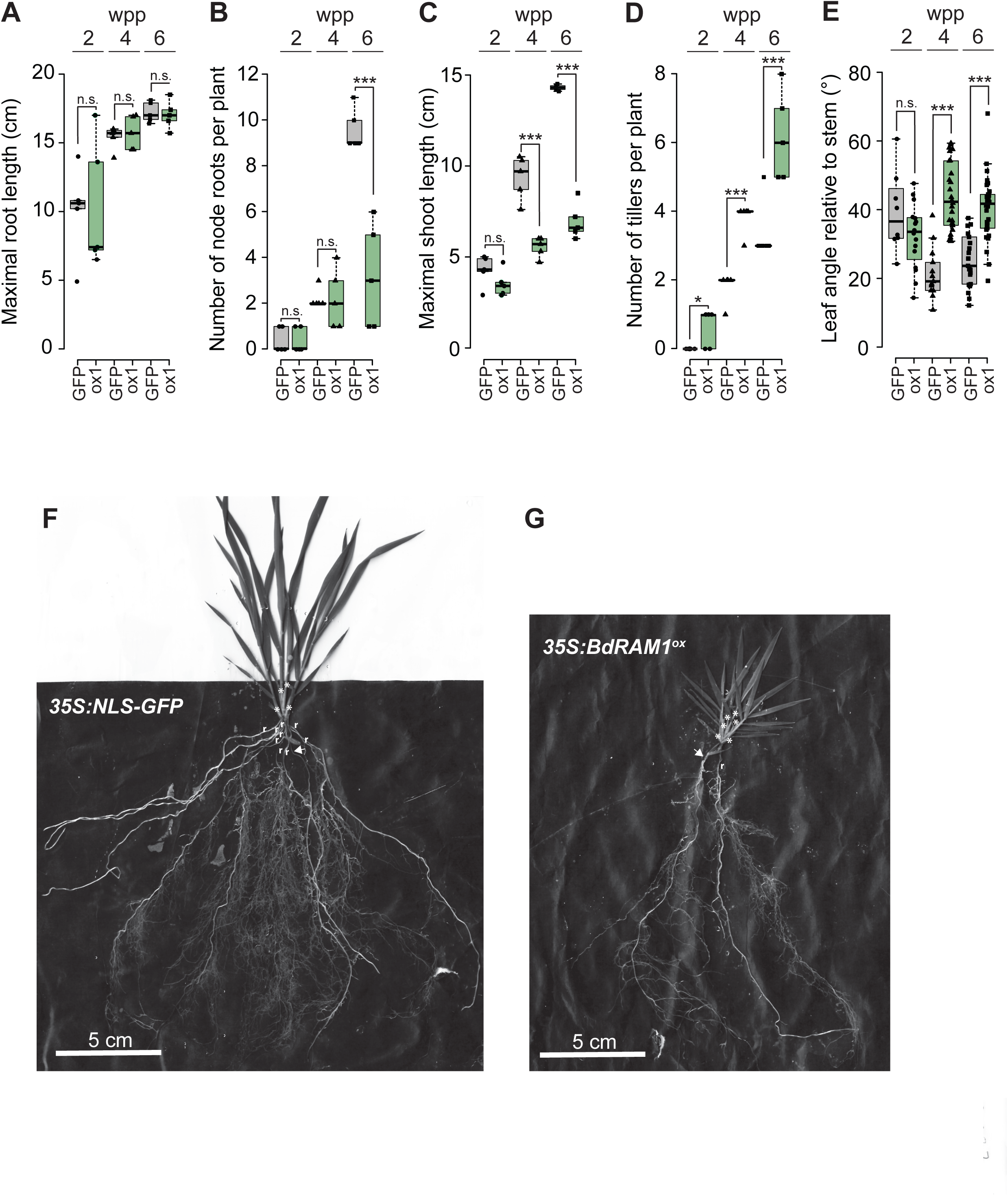
Developmental phenotypes caused by ectopic overexpression of *Bdram1* measured at 2, 4, and 6 weeks post planting (wpp). A) Maximal root system length does not differ between *355:Bdram1*^*ox*^ (line #1, “ox1”) and control plants (*355:NL5-GFP*, denoted as “GFP”). B) The collective number of coleoptile and leaf node roots is decreased in 6-week old *355:Bdram1*^*ox*^ relative to control plants. C) Overall shoot length is strongly decreased in 4- and 6-week old *355:Bdram1*^*ox*^ relative to control plants. D) Ectopic overexpression of *Bdram1* results in increased tiller numbers relative to control plants. E) The leaf angle relative to the stem is increased in in 4- and 6-week old *355:Bdram1*^*ox*^ plants relative to the control. A)-E) Box-and-whiskers plots show lower and upper quartiles, and minimum and maximum values. The horizontal bar represents the median, and the points individual measurements. Pairwise comparisons were calculated separately for each time point (Student’s t-test). Significance codes: ***p<0.001; **p<0.01; *p<0.05; n.s., not significant. F) Representative image of a plant expressing *355:NL5-GFP*. G) Representative image of a *355:Bdram1*^*ox*^ plant. F), G) Node roots are designated with “r”, and tillers are designated with “*”; arrow points to seminal root.

**Figure S5:**
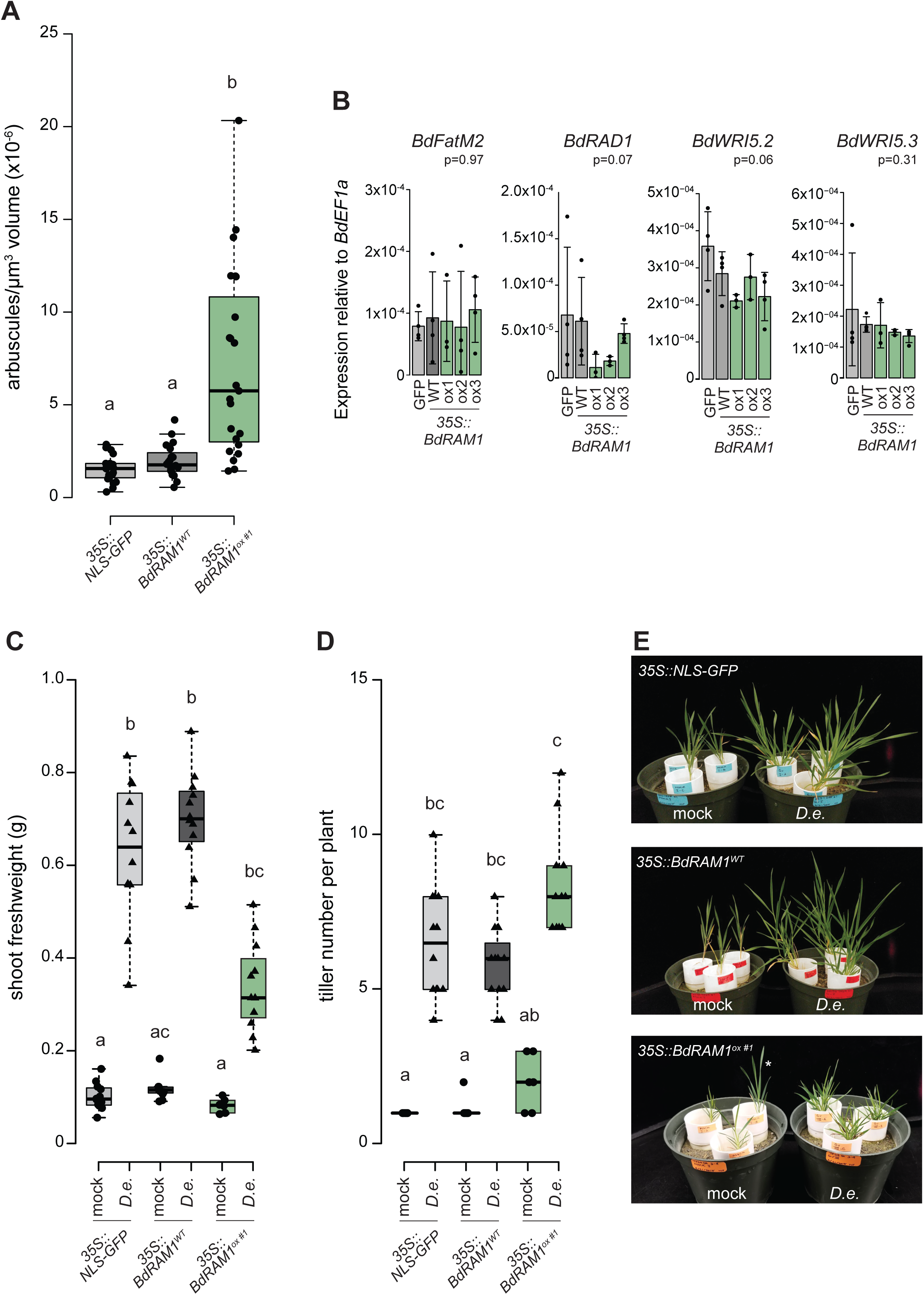
Growth response experiment and analysis of arbuscule numbers and gene expression in *355:Bdram1*^*ox*^ plants. A) Arbuscule density in a defined volume below the hyphopodium. This is a second independent experiment that shows increased arbuscule density per root volume in *355:Bdram1*^*ox*^ roots relative to *355:NL5-GFP* and *355:Bdram1*^*WT*^ control plants (compare Fig. 3D). Kruskal-Wallis test p=3.78×10^−07^. B) Gene expression levels of *BdFatM2, BdRADl, BdWR/5*.*2*, and *BdWR/5*.*3* in shoots overexpressing *Bdram1* (*355:Bdram1*^*ox*^ lines 1-3) and control roots (*355:NL5-GFP* and *355:Bdram1*^*WT*^). No significant differences were detected (p-value after ANOVA is shown in the figure). C)-E) *B. distachyon* growth response experiment: C) Shoot freshweight of 9-week old mock-inoculated (“mock”) *355:Bdram1*^*ox*^ (line 1) and *355:NL5-GFP* and *355:Bdram1*^*WT*^ control plants and plants inoculated with *D. epigaea* (“*D*.*e*.”). Kruskal-Wallis test p= 9.78×10^−11^. D) Tiller numbers of the same plants. Kruskal-Wallis test p= 8.22×10^−11^. E) Representative images of mock-inoculated and *D. epigaea*-inoculated plants at harvest (9 weeks after planting). The developmental defects of *355:Bdram1*^*ox*^ plants were so severe that we could not draw conclusions from the growth response measurements in response to AM colonization. Note: the plant marked with an asterisk is a wild-type segregant and was not included in the analysis shown in C) and D). A), C), D) Individual measurements are displayed as single points. Box plots show median as horizontal line, and upper and lower quartiles (dashed lines). Different letters denote significant differences (p<0.05) after Dunn’s posthoc test.

**Figure S6:**
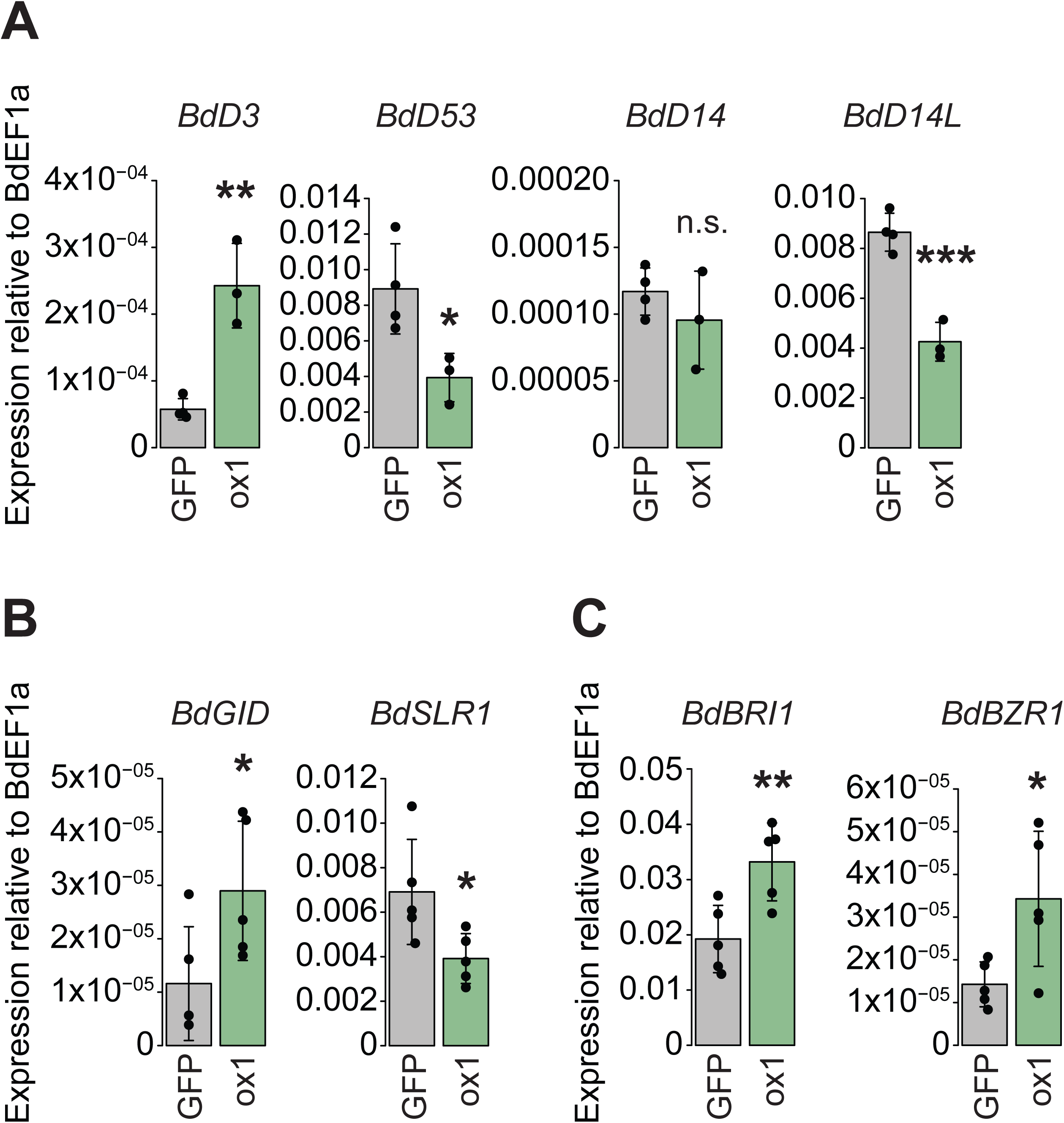
Ectopic overexpression of *Bdram1* influences expression of genes associated with strigolactone, gibberellic acid, and brassinosteroid signaling. *A) B. distachyon* orthologs of two genes involved in the strigolactone signaling pathway (*BdD3, BdD53*) are differentially regulated in non-colonized roots ectopically overexpressing *Bdram1* (*355:Bdram1*^*ox*^ line#1, denoted as “ox1”) relative to *355:NL5-GFP* (“GFP”) control roots. Gene expression of the putative strigolactone receptor *BdDl4* is not influenced by ectopic overexpression of *Bdram1*. Gene expression of the putative karrikin receptor *BdDl4L* is reduced in *Bdram1*^*ox*^ roots. B) Two genes with a putative function in Gibberelic acid signaling (*BdG/D, Bd5LRl*) are differentially regulated in *355:Bdram1*^*ox*^ roots relative to control roots. C) Two *B. distachyon* genes orthologous to known Brassinosteroid signaling genes (*BdBR/l, BdBZRl*) are induced in *355:Bdram1*^*ox*^ roots. Bar graphs show the mean, error bars the standard deviation. Single points represent individual measurements. Pairwise comparisons were estimated using the Student’s t-test. Significance codes: ***p<0.001; **p<0.01; *p<0.05; n.s., not significant.

**Figure S7:**
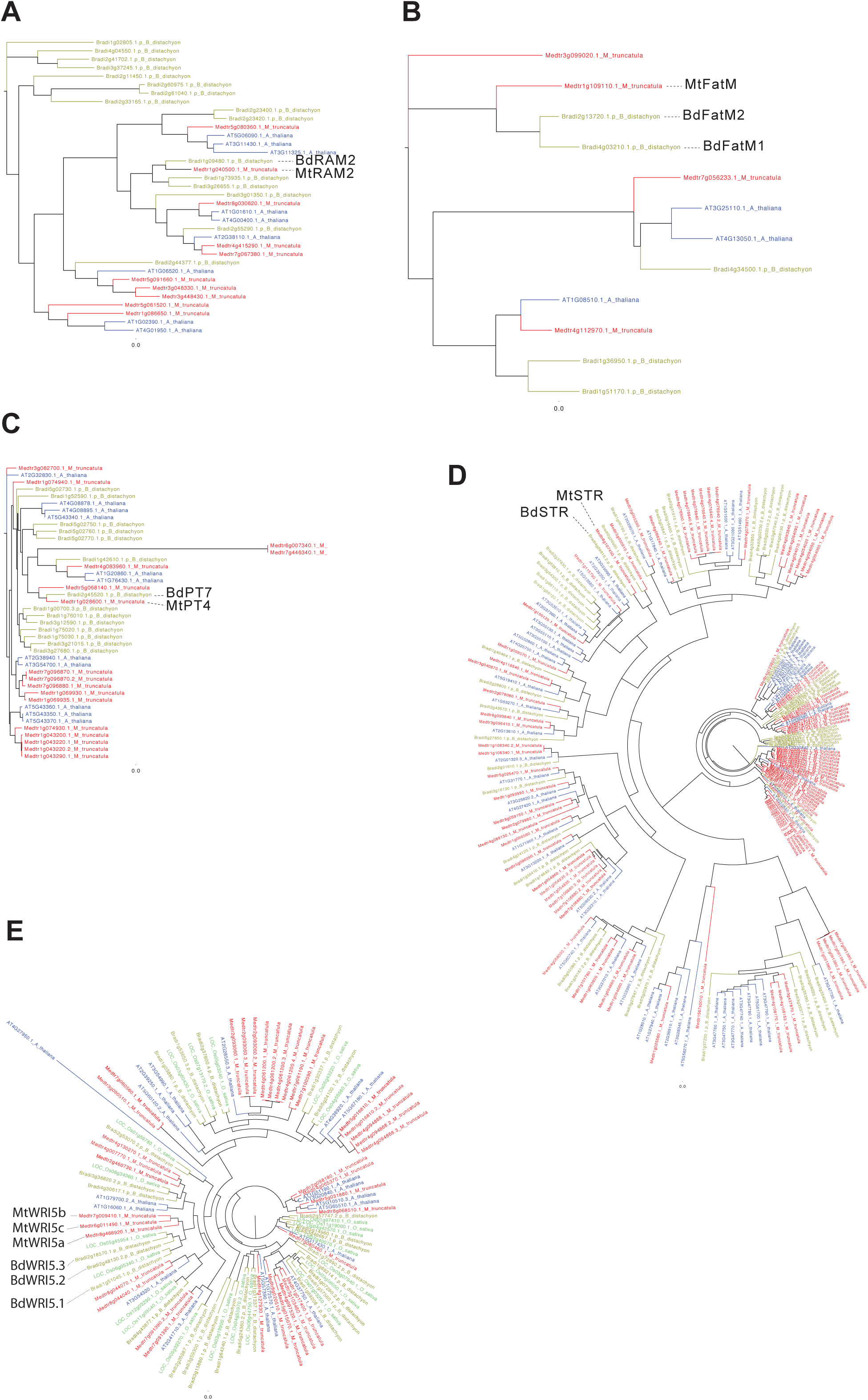
Phylogenetic trees used to identify *B. distachyon* orthologs of AM marker genes.

**Supplementary Table 1:**
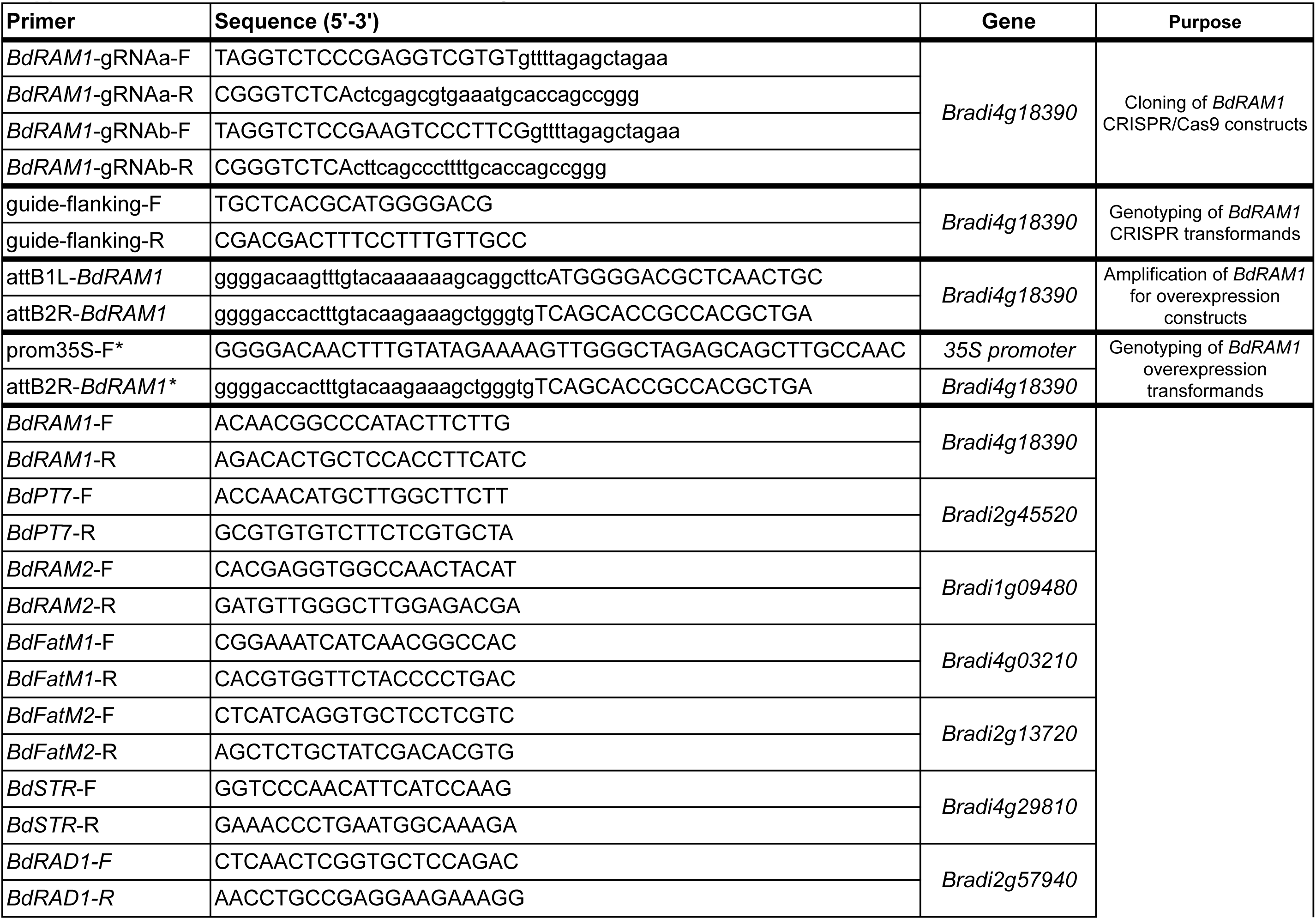
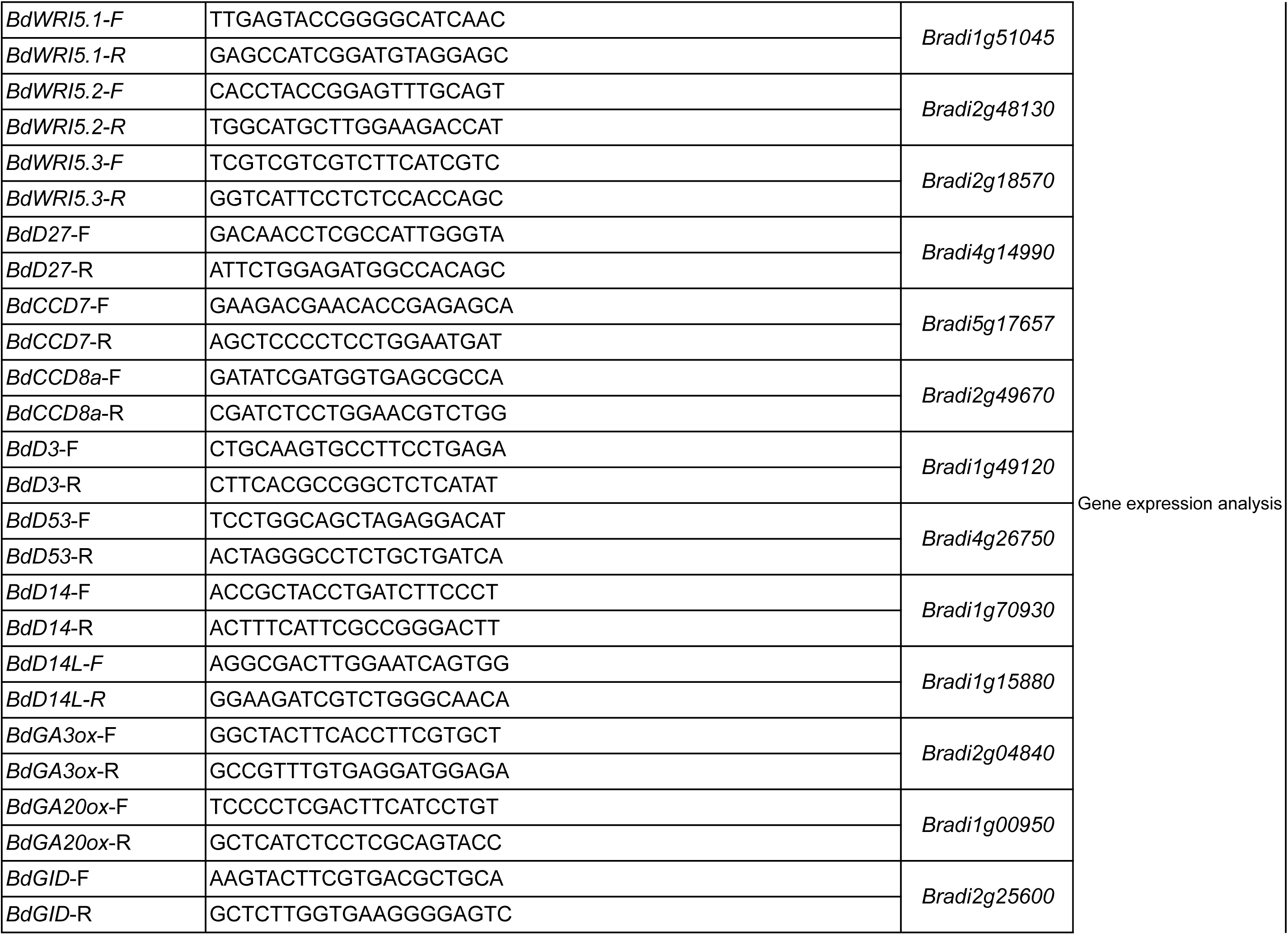
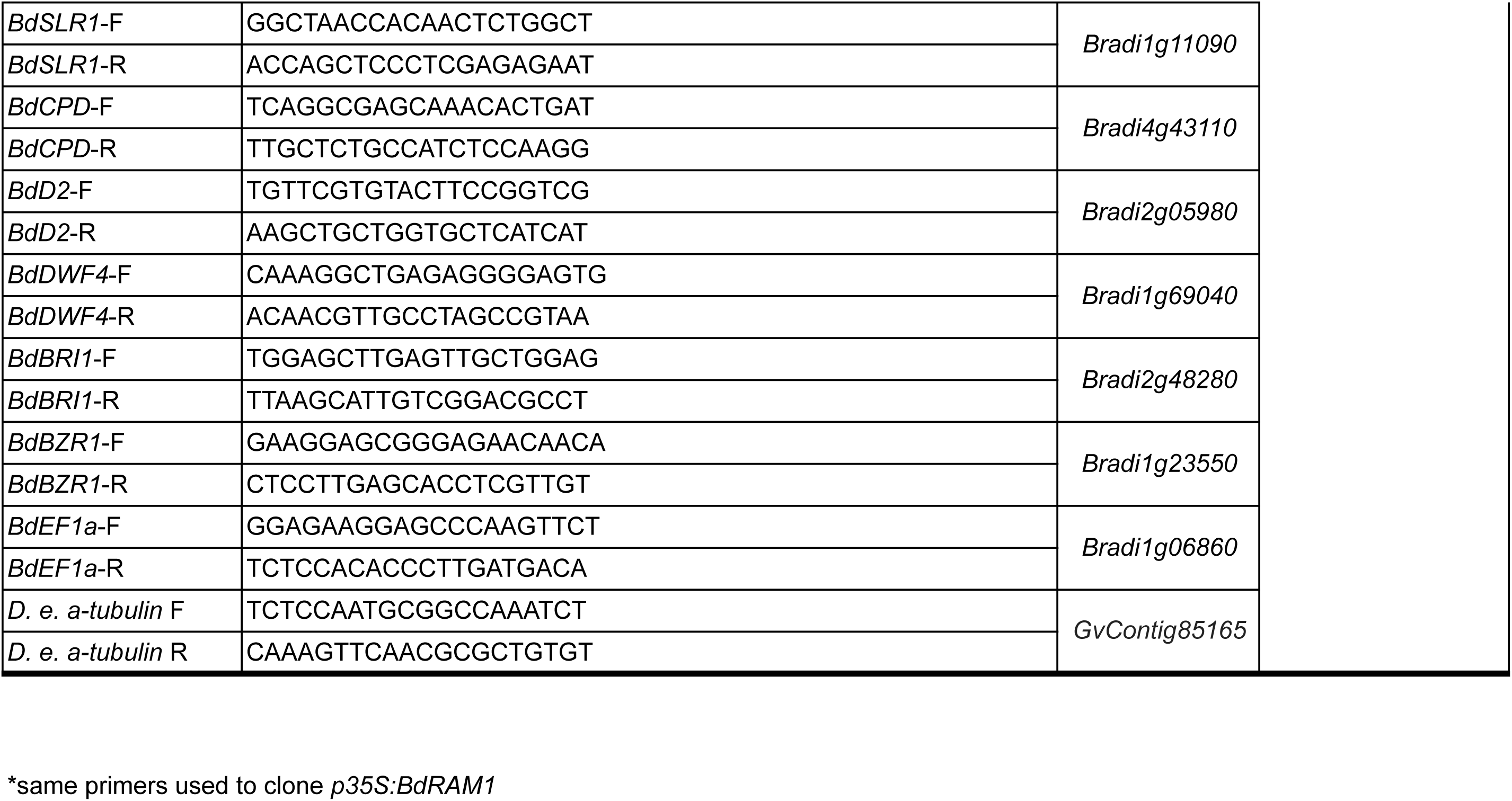
Primers used in this study.

## References

Asano K, Hirano K, Ueguchi-Tanaka M, Angeles-Shim RB, Komura T, Satoh H, Kitano H, Matsuoka M, Ashikari M (2009) Isolation and characterization of dominant dwarf mutants, Slr1-d, in rice. Molecular Genetics and Genomics 281: 223–231

Bragg JN, Anderton A, Nieu R, Vogel JP (2015) Brachypodium distachyon. In Springer, ed, Agrobacterium Protocols, Vol 1. Springer, New York, pp 17–32

Bravo A, Brands M, Wewer V, Doermann P, Harrison M J (2017) Arbuscular mycorrhiza-specific enzymes FatM and RAM2 fine tune lipid biosynthesis to promote development of arbuscular mycorrhiza. New Phytologist 214: 1631–1645

Bravo A, York T, Pumplin N, Mueller LA, Harrison MJ (2016) Genes conserved for arbuscular mycorrhizal symbiosis identified through phylogenomics. Nature Plants 2: 1–6

Cenci A, Rouard M (2017) Evolutionary Analyses of GRAS Transcription Factors in Angiosperms. Frontiers in Plant Science 8

Chen LS, Xiong GS, Cui X, Yan MX, Xu T, Qian Q, Xue YB, Li JY, Wang YH (2013) OsGRAS19 May Be a Novel Component Involved in the Brassinosteroid Signaling Pathway in Rice. Molecular Plant 6: 988–991

Corvalan C, Choe S (2017) Identification of brassinosteroid genes in Brachypodium distachyon. Bmc Plant Biology 17

Daviere JM, Achard P (2013) Gibberellin signaling in plants. Development 140: 1147–1151

Floss DS, Gomez SK, Park HJ, MacLean AM, Mueller LA, Bhattarai KK, Levesque-Tremblay V, Maldonado-Mendoza IE, Harrison M J (2017) A transcriptional program for arbuscule degeneration during AM symbiosis regulated by MYB1. Current Biology 27: 1-27

Floss DS, Levesque-Tremblay V, Park H, Harrison M J (2016) DELLA proteins regulate expression of a subset of AM symbiosis-induced genes in *Medicago truncatula*. Plant Signaling and Behaviour 11: e1162369

Floss DS, Levy JG, Levesque-Tremblay V, Pumplin N, Harrison MJ (2013) DELLA proteins regulate arbuscule formation in arbuscular mycorrhizal symbiosis. Proceedings of the National Academy of Sciences 110: doi: 10.1073/pnas.1308973110

Fonouni-Farde C, Tan S, Baudin M, Brault M, Wen JQ, Mysore KS, Niebel A, Frugier F, Diet A (2016) DELLA-mediated gibberellin signalling regulates Nod factor signalling and rhizobial infection. Nature Communications 7

Foo E, Ross JJ, Jones WT, Reid JB (2013) Plant hormones in arbuscular mycorrhizal symbioses: an emerging role for gibberellins. Annals of Botany

Gobbato E, Marsh JF, Vernie T, Wang E, Maillet F, Kim J, Miller JB, Sun J, Bano SA, Ratet P, Mysore KS, Denarie J, Schultze M, Oldroyd GED (2012) A GRAS-Type Transcription Factor with a Specific Function in Mycorrhizal Signaling. Current Biology 22: 2236–2241

Gobbato E, Wang E, Higgins G, Bano SA, Henry C, Schultze M, Oldroyd GED (2013) RAM1 and RAM2 function and expression during Arbuscular Mycorrhizal Symbiosis and Aphanomyces euteiches colonization. Plant Signaling & Behavior 8: e26049

Gomez-Roldan V, Fermas S, Brewer PB, Puech-Pages V, Dun EA, Pillot J-P, Letisse F, Matusova R, Danoun S, Portais J-C, Bouwmeester H, Becard G, Beveridge CA, Rameau C, Rochange SF (2008) Strigolactone inhibition of shoot branching. Nature 455: 189–194

Gutjahr C, Banba M, Croset V, An K, Miyao A, An G, Hirochika H, Imaizumi-Anraku H, Paszkowski U (2008) Arbuscular Mycorrhiza-Specific Signaling in Rice Transcends the Common Symbiosis Signaling Pathway. Plant Cell 20: 2989–3005

Gutjahr C, Radovanovic D, Geoffroy J, Zhang Q, Siegler H, Chiapello M, Casieri L, An K, An G, Guiderdoni E, Kumar CS, Sundaresan V, Harrison MJ, Paszkowski U (2012) The half-size ABC transporters STR1 and STR2 are indispensable for mycorrhizal arbuscule formation in rice. Plant Journal 69: 906–920

Harrison MJ, Dewbre GR, Liu J (2002) A phosphate transporter from *Medicago truncatula* involved in the acquisition of phosphate released by arbuscular mycorrhizal fungi. Plant Cell 14: 2413–2429

Heck C, Kuhn H, Heidt S, Walter S, Rieger N, Requena N (2016) Symbiotic Fungi Control Plant Root Cortex Development through the Novel GRAS Transcription Factor MIG1. Current Biology 26: 2770–2778

Hong JJ, Park YS, Bravo A, Bhattarai KK, Daniels DA, Harrison MJ (2012) Diversity of morphology and function in arbuscular mycorrhizal symbioses in Brachypodium distachyon. Planta 236: 851–865

Ishikawa S, Maekawa M, Arite T, Onishi K, Takamure I, Kyozuka J (2005) Suppression of tiller bud activity in tillering dwarf mutants of rice. Plant and Cell Physiology 46: 79–86

Itoh H, Ueguchi-Tanaka M, Sentoku N, Kitano H, Matsuoka M, Kobayashi M (2001) Cloning and functional analysis of two gibberellin 3 beta-hydroxylase genes that are differently expressed during the growth of rice. Proceedings of the National Academy of Sciences of the United States of America 98: 8909–8914

Ivanov S, Harrison MJ (2014) A set of fluorescent protein-based markers expressed from constitutive and arbuscular mycorrhiza-inducible promoters to label organelles, membranes and cytoskeletal elements in Medicago truncatula. Plant J 80: 1151–1163

Jiang Y, Wang W, Xie Q, Liu N, Liu L, Wang D, Zhang X, Yang C, Chen X, Tang D, Wang E (2017) Plants transfer lipids to sustain colonization by mutualistic mycorrhizal and parasitic fungi. Science 356: 1172–1175

Jiang YN, Xie QJ, Wang WX, Yang J, Zhang XW, Yu N, Zhou Y, Wang ET (2018) Medicago AP2-Domain Transcription Factor WRI5a Is a Master Regulator of Lipid Biosynthesis and Transfer during Mycorrhizal Symbiosis. Molecular Plant 11: 1344–1359

Jin Y, Liu H, Luo DX, Yu N, Dong WT, Wang C, Zhang XW, Dai HL, Yang J, Wang ET (2016) DELLA proteins are common components of symbiotic rhizobial and mycorrhizal signalling pathways. Nature Communications 7

Kakei Y, Mochida K, Sakurai T, Yoshida T, Shinozaki K, Shimada Y (2015) Transcriptome analysis of hormone-induced gene expression in Brachypodium distachyon. Scientific Reports 5

Karimi M, De Meyer B, Hilson P (2005) Modular cloning in plant cells. Trends in Plant Science 10: 103–105

Kobae Y, Kameoka H, Sugimura Y, Saito K, Ohtomo R, Fujiwara T, Kyozuka J (2018) Strigolactone Biosynthesis Genes of Rice are Required for the Punctual Entry of Arbuscular Mycorrhizal Fungi into the Roots. Plant and Cell Physiology 59: 544–553

Lei Y, Lu L, Liu H-Y, Li S, Xing F, Chen L-L (2014) CRISPR-P: A Web Tool for Synthetic Single-Guide RNA Design of CRISPR-System in Plants. Molecular Plant 7: 1494–1496

Lin H, Wang RX, Qian Q, Yan MX, Meng XB, Fu ZM, Yan CY, Jiang B, Su Z, Li JY, Wang YH (2009) DWARF27, an Iron-Containing Protein Required for the Biosynthesis of Strigolactones, Regulates Rice Tiller Bud Outgrowth. Plant Cell 21: 1512–1525

Liu W, Kohlen W, Lillo A, Op den Camp R, Ivanov S, Hartog M, Limpens E, Jamil M, Smaczniak C, Kaufmann K, Yang WC, Hooiveld G, Charnikhova T, Bouwmeester HJ, Bisseling T, Geurts R (2011) Strigolactone Biosynthesis in Medicago truncatula and Rice Requires the Symbiotic GRAS-Type Transcription Factors NSP1 and NSP2. Plant Cell 23: 3853–3865

Luginbuehl LH, Menard GN, Kurup S, Van Erp H, Radhakrishnan GV, Breakspear A, Oldroyd GED, Eastmond PJ (2017) Fatty acids in arbuscular mycorrhizal fungi are synthesized by the host plant. Science 356: 1175–1178

McGonigle TP, Miller MH, Evans DG, Fairchild GL, Swan JA (1990) A new method that gives an objective measure of colonization of roots by vesicular-arbuscular mycorrhizal fungi. New Phytologist 115: 495–501

Muller LM, Flokova K, Schnabel E, Sun XP, Fei ZJ, Frugoli J, Bouwmeester HJ, Harrison MJ (2019) A CLE-SUNN module regulates strigolactone content and fungal colonization in arbuscular mycorrhiza. Nature Plants 5: 933–939

Niu X, Chen SK, Li JW, Liu Y, Ji WQ, Li HF (2019) Genome-wide identification of GRAS genes in Brachypodium distachyon and functional characterization of BdSLR1 and BdSLRL1. Bmc Genomics 20

Park H, Floss DS, Levesque-Tremblay V, Bravo A, Harrison M J (2015) Hyphal branching during arbuscule development requires Reduced Arbuscular Mycorrhiza 1. Plant Physiology 169: 1–15

Paszkowski U, Kroken S, Roux C, Briggs SP (2002) Rice phosphate transporters include an evolutionarily divergent gene specifically activated in arbuscular mycorrhizal symbiosis. Proceedings of the National Academy of Sciences, USA 99: 13324–13329

Pimprikar P, Carbonnel S, Paries M, Katzer K, Klingl V, Bohmer MJ, Karl L, Floss DS, Harrison MJ, Parniske M, Gutjahr C (2016) A CCaMK-CYCLOPS-DELLA complex activates transcription of *RAM1* to regulate arbuscule branching. Current Biology 26: 987–998

Pimprikar P, Gutjahr C (2018) Transcriptional Regulation of Arbuscular Mycorrhiza Development. Plant and Cell Physiology 59: 678–695

Rich M, Schorderet M, Bapaume L, Falquet L, Morel P, Vandenbussche M, Reinhardt D (2015) The Petunia GRAS Transcription Factor ATA/RAM1 Regulates Symbiotic Gene Expression and Fungal Morphogenesis in Arbuscular Mycorrhiza. Plant Physiology DOI: 10.1104/pp.15.00310

Schindelin J, Arganda-Carreras I, Frise E, Kaynig V, Longair M, Pietzsch T, Preibisch S, Rueden C, Saalfeld S, Schmid B, Tinevez JY, White DJ, Hartenstein V, Eliceiri K, Tomancak P, Cardona A (2012) Fiji: an open-source platform for biological-image analysis. Nature Methods 9: 676–682

Seto Y, Yamaguchi S (2014) Strigolactone biosynthesis and perception. Current Opinion in Plant Biology 21: 1–6

Spielmeyer W, Ellis MH, Chandler PM (2002) Semidwarf (sd-1), “green revolution” rice, contains a defective gibberellin 20-oxidase gene. Proceedings of the National Academy of Sciences of the United States of America 99: 9043–9048

Thole V, Peraldi A, Worland B, Nicholson P, Doonan JH, Vain P (2012) T-DNA mutagenesis in Brachypodium distachyon. Journal of Experimental Botany 63: 567–576

Tong HN, Jin Y, Liu WB, Li F, Fang J, Yin YH, Qian Q, Zhu LH, Chu CC (2009) DWARF AND LOW-TILLERING, a new member of the GRAS family, plays positive roles in brassinosteroid signaling in rice. Plant Journal 58: 803–816

Umehara M, Hanada A, Yoshida S, Akiyama K, Arite T, Takeda-Kamiya N, Magome H, Kamiya Y, Shirasu K, Yoneyama K, Kyozuka J, Yamaguchi S (2008) Inhibition of shoot branching by new terpenoid plant hormones. Nature 455: 195–200

Unterholzner SJ, Rozhon W, Papacek M, Ciomas J, Lange T, Kugler KG, Mayer KF, Sieberer T, Poppenberger B (2015) Brassinosteroids Are Master Regulators of Gibberellin Biosynthesis in Arabidopsis. Plant Cell 27: 2261–2272

Watts-Williams SJ, Emmett BD, Levesque-Tremblay V, MacLean AM, Sun X, Satterlee JW, Fei Z, Harrison MJ (2019) Diverse Sorghum bicolor accessions show marked variation in growth and transcriptional responses to arbuscular mycorrhizal fungi. Plant Cell and Environment 42: 1758–1774

Xie K, Minkenberg B, Yang Y (2015) Boosting CRISPR/Cas9 multiplex editing capability with the endogenous tRNA-processing system. Proceedings of the National Academy of Sciences 112: 3570–3575

Xue L, Cui H, Buer B, Vijayakumar V, Delaux P-M, Junkermann S, Bucher M (2015) Network of GRAS transcription factors involved in the control of arbuscule development in *Lotus japonicus*. Plant Physiology 167: 854–871

Yu N, Luo D, Zhang X, Liu J, Wang W, Jin Y, Dong W, Liu J, Liu H, Yang W, Zeng L, Li Q, He Z, Oldroyd GED, Wang E (2014) A DELLA protein complex controls the arbuscular mycorrhizal symbiosis in plants. Cell Res 24: 130–133

Zhang Q, Blaylock LA, Harrison M J (2010) Two *Medicago truncatula* half-ABC transporters are essential for arbuscule development in arbuscular mycorrhizal symbiosis. Plant Cell 22: 1483–1497

## References

Bravo, A., York, T., Pumplin, N., Mueller, L.A., and Harrison, M.J. (2016). Genes conserved for arbuscular mycorrhizal symbiosis identified through phylogenomics. Nature Plants 2, 1–6.

Chen, L.S., Xiong, G.S., Cui, X., Yan, M.X., Xu, T., Qian, Q., Xue, Y.B., Li, J.Y., and Wang, Y.H. (2013). OsGRAS19 May Be a Novel Component Involved in the Brassinosteroid Signaling Pathway in Rice. Mol. Plant 6, 988–991.

Floss, D.S., Levesque-Tremblay, V., Park, H., and Harrison M, J. (2016). DELLA proteins regulate expression of a subset of AM symbiosis-induced genes in *Medicago truncatula*. Plant Signaling and Behaviour 11, e1162369.

Floss, D.S., Levy, J.G., Levesque-Tremblay, V., Pumplin, N., and Harrison, M.J. (2013). DELLA proteins regulate arbuscule formation in arbuscular mycorrhizal symbiosis. Proceedings of the National Academy of Sciences 110, doi: 10.1073/pnas.1308973110.

Floss, D.S., Gomez, S.K., Park, H.J., MacLean, A.M., Mueller, L.A., Bhattarai, K.K., Levesque-Tremblay, V., Maldonado-Mendoza, I.E., and Harrison M, J. (2017). A transcriptional program for arbuscule degeneration during AM symbiosis regulated by MYB1. Current Biology 27, 1–27.

Liu, W., Kohlen, W., Lillo, A., Op den Camp, R., Ivanov, S., Hartog, M., Limpens, E., Jamil, M., Smaczniak, C., Kaufmann, K., Yang, W.C., Hooiveld, G., Charnikhova, T., Bouwmeester, H.J., Bisseling, T., and Geurts, R. (2011). Strigolactone Biosynthesis in Medicago truncatula and Rice Requires the Symbiotic GRAS-Type Transcription Factors NSP1 and NSP2. Plant Cell 23, 3853–3865.

McGonigle, T.P., Miller, M.H., Evans, D.G., Fairchild, G.L., and Swan, J.A. (1990). A new method that gives an objective measure of colonization of roots by vesicular-arbuscular mycorrhizal fungi. New Phytol. 115, 495–501.

Park, H., Floss, D.S., Levesque-Tremblay, V., Bravo, A., and Harrison M, J. (2015). Hyphal branching during arbuscule development requires Reduced Arbuscular Mycorrhiza 1. Plant Physiol. 169, 1–15.

Tong, H.N., Jin, Y., Liu, W.B., Li, F., Fang, J., Yin, Y.H., Qian, Q., Zhu, L.H., and Chu, C.C. (2009). DWARF AND LOW-TILLERING, a new member of the GRAS family, plays positive roles in brassinosteroid signaling in rice. Plant J. 58, 803–816.

Yu, N., Luo, D., Zhang, X., Liu, J., Wang, W., Jin, Y., Dong, W., Liu, J., Liu, H., Yang, W., Zeng, L., Li, Q., He, Z., Oldroyd, G.E.D., and Wang, E. (2014). A DELLA protein complex controls the arbuscular mycorrhizal symbiosis in plants. Cell Res. 24, 130–133.

